# Quantitative and sensitive neuroblastoma minimal residual disease detection using extrachromosomal DNA (ecDNA) breakpoints

**DOI:** 10.64898/2026.09.06.749680

**Authors:** Annabell Szymansky, Falk Hertwig, Annika Winkler, Rocío Chamorro González, Konstantin Helmsauer, Anna-Marie Schroeer, Melanie J. Witthauer, Eliz Karaman, Marco Lodrini, Jasmin Wünschel, Nadine Wittstruck, Steffen Fuchs, Jessica Theissen, Barbara Hero, Richard Koche, Malwine J. Barz, Lukas C. Heukamp, Kathy Astrahantseff, Karin Schmelz, Patrick Hundsdoerfer, Joern Toedling, Angelika Eggert, Cornelia Eckert, Bastiaan Spanjaard, Hedwig E. Deubzer, Matthias Fischer, Anton G. Henssen, Johannes H. Schulte

**Author notes:** Corresponding author: Annabell Szymansky, PhD, Charité – Universitätsmedizin Berlin, Department of Pediatric Oncology and Hematology, Augustenburger Platz 1, 13353 Berlin, Germany, T. These authors contributed equally to this work (Hedwig E. Deubzer, Matthias Fischer, Anton G. Henssen, Johannes H. Schulte).

## Abstract

Sensitive detection of minimal residual disease (MRD) remains a major unmet need in high-risk neuroblastoma. *MYCN* amplification, a hallmark of high-risk disease, typically occurs on extrachromosomal DNA (ecDNA), but the potential of ecDNA-associated genomic rearrangements for individualized MRD monitoring has not been fully exploited. Here, we applied neuroblastoma-specific hybrid capture-based panel sequencing to identify patient-unique breakpoints within *MYCN* amplicons, and used Circle-seq and Nanopore sequencing to resolve the extrachromosomal amplicon structure in representative samples. Analysis of 8 neuroblastoma cell lines and 22 primary tumors identified 69 tumor-specific breakpoints. Those selected for assay development were validated by breakpoint-specific PCR and Sanger sequencing. Breakpoints detected in primary tumors remained detectable at relapse, supporting their stability as MRD markers. Breakpoint-specific real-time quantitative PCR and droplet digital PCR detected these junctions in bone marrow aspirates with high specificity and reached sensitivities down to a tumor DNA fraction of 10⁻⁶. We applied this approach to 53 serial bone marrow aspirates from 14 patients with high-risk neuroblastoma to monitor MRD dynamics, resolving treatment response and molecular persistence. In six samples, breakpoint-positive DNA was detected in bone marrow that was negative by conventional cytology and immunocytology, highlighting the added value of molecular monitoring. Together, these findings establish ecDNA breakpoint-based detection as a strategy for MRD assessment in neuroblastoma, that is, in principle, applicable to any ecDNA-amplified oncogene.

**Single Sentence Summary:** Breakpoints in extrachromosomal circular DNA, a substrate of oncogene amplification in cancer, can serve as diagnostic markers of minimal residual disease.

## Introduction

Neuroblastoma represents the most common extracranial solid malignancy of childhood, presenting an exceptionally heterogeneous clinical course that ranges from spontaneous regression in infants to high-risk, aggressively metastatic disease(1). Systemic dissemination is initially diagnosed in approximately 50% of patients, with the bone marrow serving as a frequent metastatic site. Disseminated neuroblastoma cells in the bone marrow often survive initial systemic multimodal therapy to persist as minimal residual disease (MRD). The persistence of neuroblastoma cells in the bone marrow is a predictor of overall survival and correlates with relapse(2,3).

Immunocytology of bone marrow smears and histopathological evaluation of core trephine biopsies have remained the primary baseline investigations to define disease stage and assess therapeutic response(4–6). However, these conventional standard-of-care diagnostics lack sufficient analytical sensitivity, frequently leaving MRD undetected(7). Multi-modal high-sensitivity assays such as multiparameter flow cytometry(8), automated immunofluorescence combined with fluorescence in situ hybridization (FISH; 9) and reverse transcription quantitative PCR (RT-qPCR; 10) have consequently been developed to lower the limit of detection. Despite these technological advances, current diagnostic paradigms have remained constrained by anatomical, technical and biological bottlenecks. The highly uneven focal distribution of neuroblastoma cells in the bone marrow creates spatial heterogeneity. This physical limitation is exacerbated by the cytotoxic effects of intensive induction chemotherapy, which reduces bone marrow cellularity and thus limits cellular representation for morphological analyses. Technically, transcript-based assays rely on mRNA, an inherently less stable analyte than genomic DNA. Biologically, antibody-dependent detection methods can be compromised by therapeutic antigenic escape, wherein neuroblastoma cells show a complete or partial absence of the surface disialoganglioside GD2(11), and can result in false-negative readouts despite persistent active disease. Further, neuroblastoma cells can undergo phenotypic plasticity(12,13), transitioning from an adrenergic state to a chemotherapy-resistant mesenchymal state in which GD2 expression is reduced and anti-GD2 antibody activity is impaired(14). Conventional RT-qPCR marker panels target adrenergic-associated transcripts such as tyrosine hydroxylase and *PHOX2B*(15). Mesenchymal tumor cell populations thus escape molecular detection, representing a diagnostic blind spot in MRD monitoring(16).

*MYCN* amplification is present in approximately 20% of neuroblastomas overall(5) and in about 30–40% of high-risk tumors(17) and remains one of the most important molecular markers associated with poor prognosis. In more than 90% of cases, *MYCN* amplification occurs on extrachromosomal DNA (ecDNA, historically termed double minutes; 18,19). Recent studies have revealed an unanticipated abundance of ecDNA across cancer types(20). These elements contribute to intratumoral heterogeneity, tumor evolution, therapy resistance and adverse clinical outcome(21–23). Because ecDNA arises through structural rearrangement, each amplicon carries junction sequences that are unique to the individual tumor(19,20). We hypothesized that these highly specific *MYCN* breakpoints in neuroblastoma represent substrates for MRD detection, analogous to the genomic fusion and antigen-receptor rearrangements established in hematologic malignancies(24,25). Using hybrid capture-based panel sequencing, we identified patient-specific *MYCN* breakpoint sequences and resolved their extrachromosomal localization by Circle-seq and Nanopore sequencing. These breakpoints were then used to design personalized real-time quantitative PCR (RQ-PCR) and droplet digital PCR (ddPCR) assays for longitudinal MRD assessment in the bone marrow.

## Results

### Hybrid capture-based panel sequencing identifies patient-specific *MYCN* amplicon breakpoints

To detect *MYCN* ecDNA in neuroblastoma, we applied fluorescence *in situ* hybridization (FISH) to interphase cells in 8 neuroblastoma cell lines and 22 primary high-risk neuroblastomas. Even though metaphase spreads are the gold standard to distinguish ecDNA from homogeneously staining regions (HSR), amplification status can be estimated based on the fluorescence patterns of FISH stained interphase cells(23). FISH patterns indicated ecDNA amplification of *MYCN* amplified on ecDNA in 3 of 8 cell lines (LAN-5, TR-14 and CHP-212; Table 1). FISH was evaluable in 12 of the 22 tumors, of which 9 showed a dispersed signal pattern consistent with ecDNA and 3 showed clustered signals indicating intrachromosomal amplification as HSR (Table 2, Figure 1a and Supplemental Figure S1, (26)). To detect breakpoints on *MYCN* amplicons, we analyzed whole-genome sequences from 14 primary *MYCN*-amplified neuroblastomas(20,27,28). Within two megabases around *MYCN*, 376 breakpoints were identified (Supplemental Figure S2a). In line with our previous reports(19,20), the majority of breakpoints were located within the *MYCN* gene, upstream of *MYCN* in the intergenic area between *MYCN* and *DDX1*(29), or downstream of *MYCN* in the intergenic region between *MYCN* and *FAM49A*(30). Using this 2Mb region as the target region for *MYCN* breakpoint detection, we designed hybrid capture probes tiling this region to enable hybrid capture-based sequencing (*MYCN* amplicon sequencing assay 31; Supplemental Figure S2b). We applied this assay to the same 8 neuroblastoma cell lines and 22 primary high-risk neuroblastomas, which resulted in a mean coverage of 435x to 2075x within the 2Mb target region around *MYCN* (Supplemental Tables S1 and S2). Discordant sequencing read pairs enabled the detection of 22 breakpoints in cell lines (mean: 2.75 per cell line, Table 1) and 47 breakpoints in tumor samples (mean: 2.14 per tumor; Table 2), three of them detected only in relapse biopsies (see below). Fewer breakpoints per tumor were detected than in the whole-genome sequencing (WGS) datasets used to define the target region, reflecting both the different cohorts analyzed and the requirement that at least one breakpoint partner map within the 2 Mb hybrid-capture region. The absolute *MYCN* copy number did not significantly correlate with the number of breakpoints detected in cell lines or tumors (Spearman correlation, p=0.25 and p=0.58, respectively), nor was the copy number significantly different between samples with ecDNA or HSR-integrated amplicons (Mann-Whitney U test, p=0.39 and p=0.86, respectively). Of all 69 breakpoints identified, only 4 occurred within the *MYCN* gene itself (Supplemental Tables S1 and S2), of which 2 were in the intron between exons 2 and 3 and the other 2 in exons 2 and 3. Of the 65 breakpoints located adjacent to *MYCN*, 9 occurred within the *NBAS* gene, 4 occurred in other genes and 52 occurred in intergenic regions (Supplemental Figure S2b). All 13 breakpoints located within genes other than *MYCN* were detected in intronic and not protein-coding regions. Patient #9, whose primary tumor sample was selected for orthogonal validation, showed *MYCN* amplified on ecDNA as confirmed by FISH (Figure 1a). To validate that our neuroblastoma hybrid-capture sequencing panel(31) detected breakpoints on ecDNAs, our results were compared with Circle-seq, a method allowing enrichment and sequencing of ecDNA(20,32), performed on the tumor sample from patient #9 (Supplemental Figure S2c). Moreover, the reconstruction of the ecDNAs originating from the linear chromosome was supported by Nanopore DNA sequencing (Figure 1b,19). Importantly, a major breakpoint clone detected using Circle-seq (Supplemental Figure S2c) in the tumor sample from patient #9 confirmed the location on ecDNA, consistent with the same breakpoint detected by our capture-based sequencing assay (Supplemental Figure S2d). This breakpoint was further orthogonally validated using a breakpoint-specific PCR followed by Sanger sequencing (Figure 1c-1e). Thus, hybrid capture-based sequencing identifies patient-specific breakpoints on tumor-specific ecDNA.

**Figure 1.**
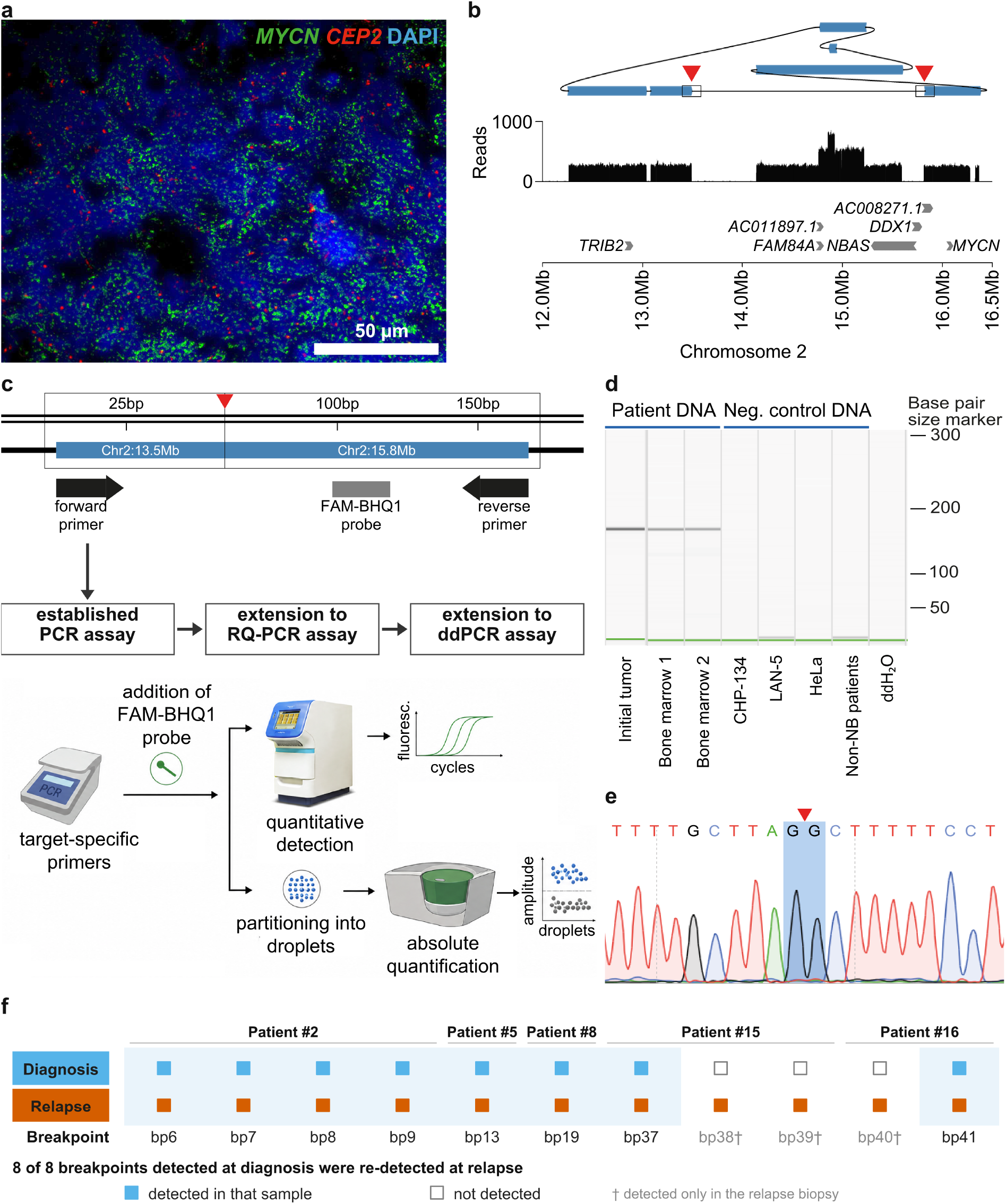
Patient-specific identification and validation of *MYCN* amplicon breakpoints. Panels (a) to (e) show patient #9. (a) Interphase FISH. The dispersed *MYCN* signal indicates amplification on extrachromosomal DNA (ecDNA). Scale bar, 50 µm. (b) Amplicon reconstruction supported by Nanopore sequencing, above hybrid capture read coverage and the gene track. (c) Breakpoint-spanning PCR assay for the chr2:13.5 Mb to chr2:15.8 Mb junction and its stepwise extension to RQ-PCR and ddPCR. The FAM-BHQ1 probe lies within the patient-specific amplicon segment without spanning the junction and is used only for the quantitative assays. (d) PCR products separated by capillary gel electrophoresis, with patient and negative control DNA as indicated. (e) Sanger sequencing of the same product, confirming the junction sequence. Red triangles in (b), (c) and (e) mark the junction. (f) Retention of *MYCN* amplicon breakpoints from the initial biopsy to relapse in the five patients with paired biopsies. Filled squares, detected by hybrid capture sequencing. Open squares, not detected. The tinted band encloses the breakpoints detected at both time points. All 8 breakpoints detected at diagnosis were re-detected at relapse. Three further breakpoints (†) were detected only in the relapse biopsy.

**Table 1:** Breakpoints detected in the *MYCN* locus in neuroblastoma cell lines.

| Name of cell line | Copy number<br>*MYCN* | *MYCN* as HSR<br>or DM | *MYCN* associated<br>breakpoints |
| --- | --- | --- | --- |
| IMR-5 | 49.90 | HSR | 2 |
| SK-N-BE | 68.06 | HSR | 3 |
| LAN-5 | 59.50 | DM | 2 |
| NGP | 56.93 | HSR | 3 |
| KELLY | 88.00 | HSR | 3 |
| TR-14 | 57.00 | DM | 3 |
| SK-N-DZ | 43.20 | HSR | 5 |
| CHP-212 | 101.00 | DM | 1 |

Breakpoints and *MYCN* copy numbers were detected by hybrid capture-based panel sequencing. Breakpoints selected for MRD assay development were validated by breakpoint-specific PCR and Sanger sequencing.

**Tablec 2:**
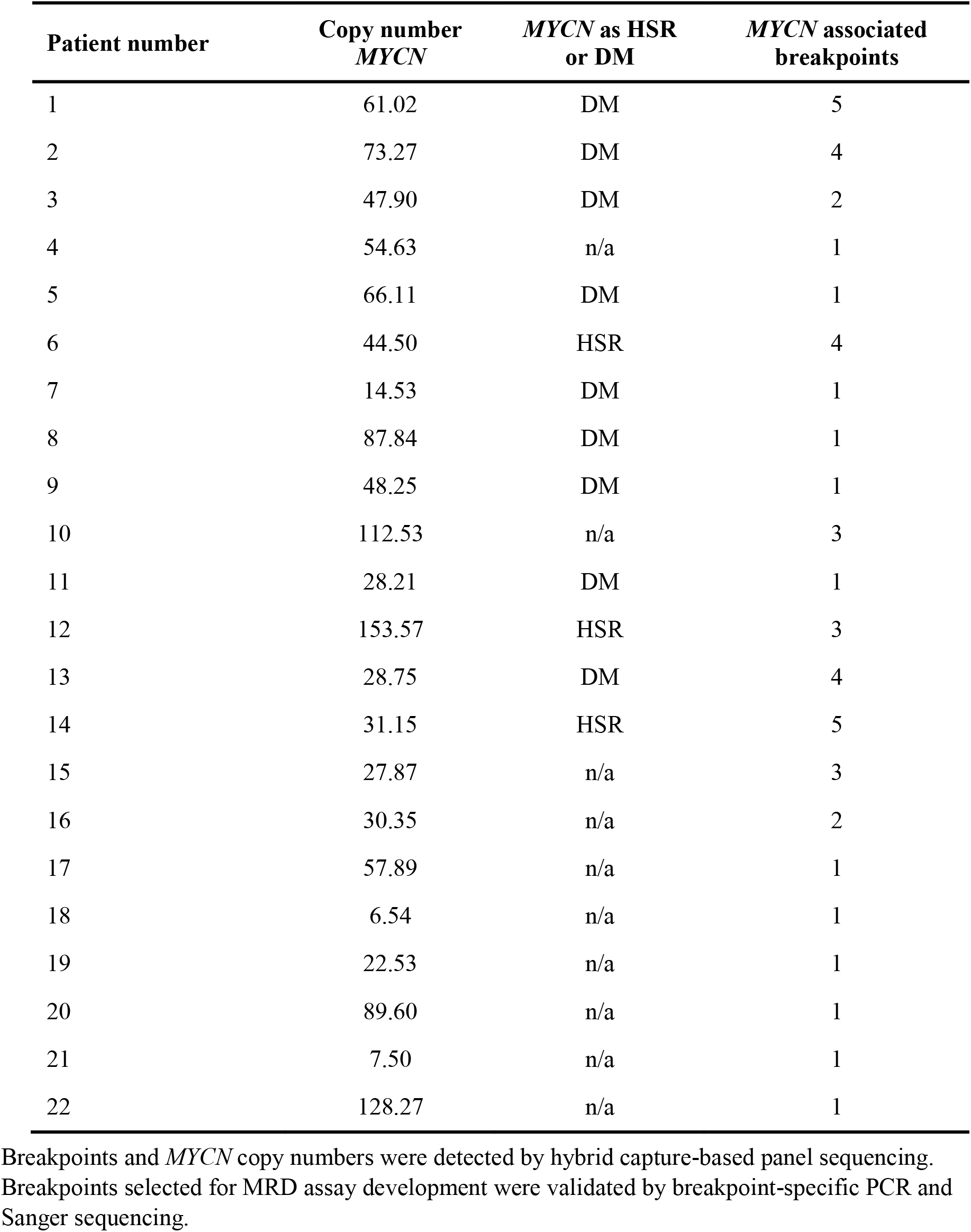
Breakpoints detected in the *MYCN* locus in primary neuroblastoma samples.

### *MYCN* amplicon breakpoints detected in primary tumors are present at relapse

The presence of breakpoints throughout disease is a prerequisite for their use as MRD markers. To verify that breakpoints detected in primary tumors persisted over the disease course and particularly at relapse, we used our capture-based sequencing assay to map *MYCN* breakpoints in paired tumor biopsies from 5 patients at diagnosis or during therapy compared to biopsies at relapse (patients 2,5,8,15 and 16; Table 2). *MYCN* FISH performed on samples from 3 patients (#2, #5, #8) indicated *MYCN* amplification on ecDNA (Supplemental Figure S1). Each of the 5 patients carried at least one *MYCN* amplicon breakpoint in the initial biopsy, and our capture-based sequencing assays detected all 8 of these breakpoints identified in primary tumors also in the matched relapsed tumor samples (Supplemental Figure S3), which was validated using breakpoint-specific PCR followed by Sanger sequencing in all cases (Figure 1f, Supplemental Table S3 and Supplemental Figure S6). Three further breakpoints were detected in relapse material only, two in patient #15 (bp38 and bp39) and one in patient #16 (bp40). This is consistent with ongoing amplicon evolution, although breakpoints present subclonally below the detection limit of the initial biopsy cannot be excluded. Thus, the *MYCN* amplicon breakpoints present in primary tumors persist through the course of disease until relapse occurs, fulfilling the prerequisites to serve as MRD markers.

### Breakpoint-specific RQ-PCR and ddPCR assays detect *MYCN* amplicon breakpoints with high sensitivity and specificity

We next sought to build a quantitative assay for breakpoint detection using TR-14, a cell line containing *MYCN* on ecDNA (Supplemental Figure S1). Breakpoints detected by our capture-based sequencing assay in TR-14 were also detected using WGS and Circle-seq (Figure 2a top, Supplemental Figure S4), confirming their location on ecDNA and enabling reconstruction of the full ecDNA structure, from which a *MYCN* breakpoint-spanning PCR was designed (Figure 2a bottom). Breakpoints selected for MRD assay development were confirmed using breakpoint-specific PCR followed by Sanger sequencing (Supplemental Figures S5 and S6), establishing breakpoint-specific PCR as a reliable and generalizable readout that could be applied across our full cohort of cell lines and patient samples. Following the quality guidelines set by the *EuroMRD* Consortium for clinically used leukemia MRD assays(33–35), we developed quantitative PCR assays for 9 breakpoints in 7 cell lines and 23 breakpoints in 22 primary tumors (Supplemental Tables S4 and S5). Each assay combined primers flanking the respective breakpoint with a dual-labeled fluorescent probe, enabling breakpoint detection through unquenching of the FAM fluorophore (Supplemental Tables S6 and S7); all assays could be run as either RQ-PCR (Figure 2b) or ddPCR (Figure 2c), the latter previously reported to offer greater precision(36) and reproducibility(37) alongside compatibility with smaller sample quantities (38). While RQ-PCR conditions were individually optimized for each assay, ddPCR was performed under standard conditions without assay-specific optimization. Using the TR-14 and their limited dilution as a proof of concept, both RQ-PCR and ddPCR achieved sensitivities of up to 10⁻⁶. Direct comparison of the two methods in TR-14 showed concordant quantification across the dilution series, with ddPCR copies/µl tracking closely with RQ-PCR Ct values across the range of MRD concentrations tested (Figure 2d); as expected and consistent with previous reports(38), ddPCR reached higher sensitivity at lower copy-number dilutions than RQ-PCR. Across the 9 cell-line breakpoints, sensitivity ranged from 10⁻⁴ to 10⁻⁶ for both RQ-PCR and ddPCR, with no significant difference between the two methods (Wilcoxon signed-rank test, p = 0.125; Figure 2e, Supplemental Table S4). Across the 23 tumor breakpoints, however, sensitivity ranged from 10⁻³ to 10⁻⁶ for both RQ-PCR and ddPCR, with ddPCR showing significantly greater sensitivity than RQ-PCR (Wilcoxon signed-rank test, p = 0.02, Figure 2f, Supplemental Table S5). No breakpoint-specific signal was detected in *MYCN*-amplified non-target neuroblastoma DNA, non-neuroblastoma DNA, pooled DNA from ten healthy donors or no-template controls run alongside every assay (e.g. Figure 2b, 2c). Together, this establishes *MYCN* breakpoint-specific RQ-PCR and ddPCR for patient-specific MRD detection.

**Figure 2.**
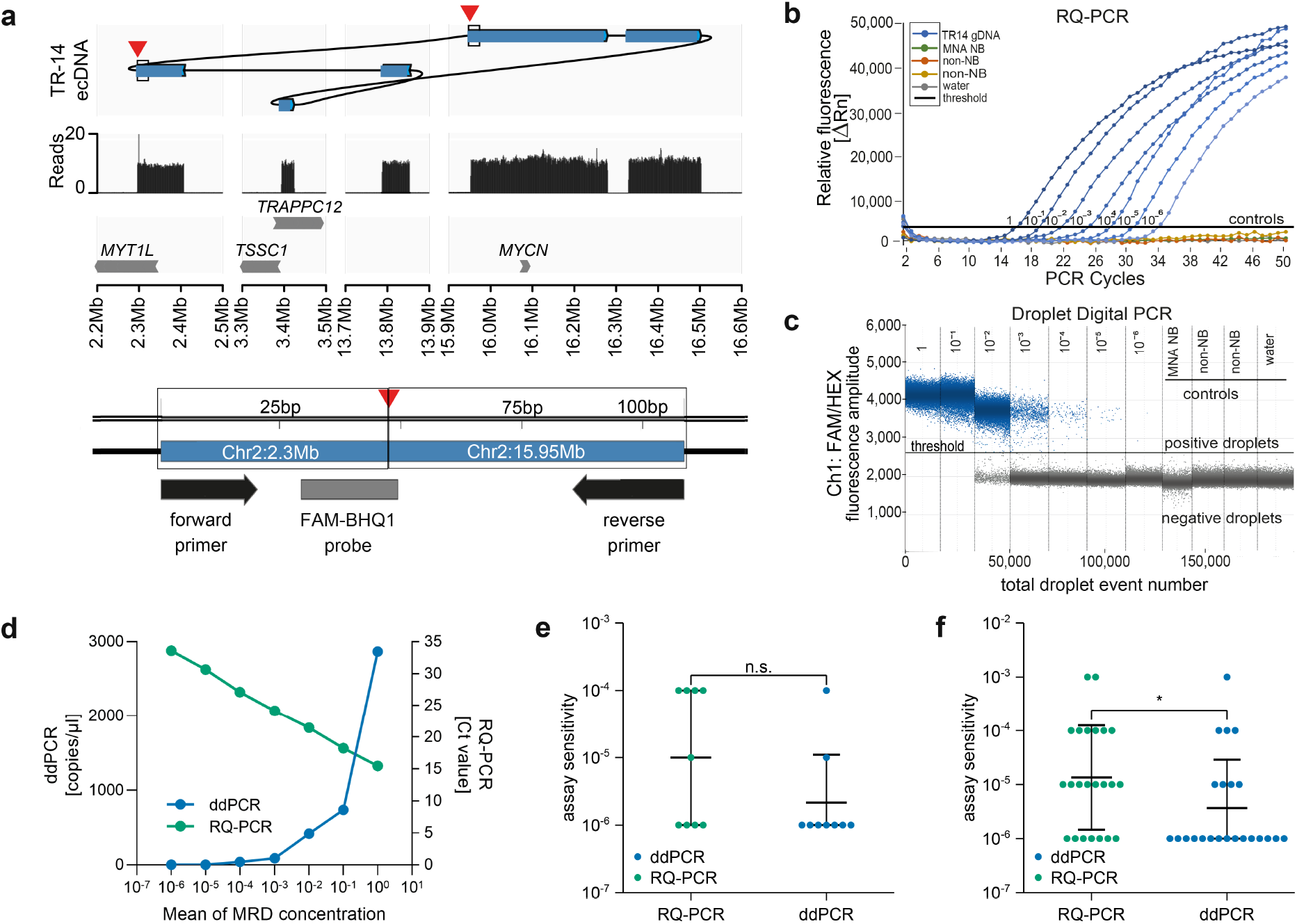
Development and performance of breakpoint-specific RQ-PCR and ddPCR assays. (a) Top: the ecDNA carrying *MYCN* in the cell line TR-14, reconstructed from whole-genome and Circle-seq data, read coverage and the gene track. Bottom: the corresponding breakpoint-spanning PCR assay. (b, c) RQ-PCR (b) and ddPCR (c) for TR-14 across a tenfold dilution series in DNA from healthy donors, with the controls indicated. (d) The two platforms compared across the same series. (e, f) Assay sensitivity of each platform for the 9 breakpoints established in 7 cell lines (e) and the 23 breakpoints established in 22 primary tumors (f). Each point is one assay. Bars give the geometric mean and geometric standard deviation. The dilution series stopped at 10⁻⁶, so assays still positive there are plotted at that value and bars reaching below it are truncated. Wilcoxon signed-rank test, n.s., p = 0.125 in (e) and *, p = 0.02 in (f).

### Application of *MYCN* breakpoint MRD assays to serial bone marrow aspirates reveals dynamic MRD in patients with neuroblastoma

To assess the clinical applicability of our assays, we developed personalized MRD assays for 14 patients for whom neuroblastoma hybrid-capture sequencing had been performed as part of the clinical diagnostics(31) and assay sensitivity had been benchmarked (Figure 3, Supplemental Figure S7, Supplemental Table S8). The assays were applied to 53 serial bone marrow aspirates collected at diagnosis and throughout treatment from patients treated according to the German high-risk neuroblastoma guidelines(39; patients #9–22 in Table 2; clinical annotation in Supplemental Table S9). Breakpoint-specific RQ-PCR and ddPCR assays yielded highly concordant results and enabled quantitative tracking of neuroblastoma cell burden over the course of therapy. Representative longitudinal profiles from six patients (patients #9, #20, #10, #19, #13, #16) demonstrated distinct patterns of molecular disease clearance (Figure 3a–f). Patients #9 and #20 achieved molecular remission during induction therapy. Patient #9 showed elevated molecular disease levels at diagnosis that rapidly declined below the assay detection limit following treatment initiation and remained undetectable thereafter (Figure 3a). Similarly, patient #20 achieved molecular remission shortly after the first treatment cycle, with breakpoint signals remaining undetectable throughout follow-up despite later clinical suspicion of relapse (Figure 3b). In contrast, patients #10 and #19 remained MRD positive during early treatment and achieved molecular remission only prior to stem cell transplantation (Figure 3c, d). Both patients exhibited substantial molecular disease burden at diagnosis, followed by a gradual decline in breakpoint levels until they became undetectable before transplantation. Patients #13 and #16 showed a different response pattern, with molecular remission achieved only after anti-GD2-based or anti-ALK-based targeted therapy and maintained during follow-up (Figure 3e, f). Across all six patients, breakpoint detection was highly concordant with bone marrow involvement identified by cytology or GD2 immunocytology. Notably, breakpoint signals were additionally detected in six samples that were negative in conventional methods, showing that personalized *MYCN* breakpoint assays provide molecular information beyond standard bone marrow diagnostics. Collectively, breakpoint-based molecular monitoring provided a continuous and quantitative assessment of treatment response across different phases of therapy. The assays revealed marked interpatient variability in disease clearance kinetics, ranging from rapid elimination of breakpoint-positive cells to gradual declines over multiple treatment cycles. Personalized *MYCN* breakpoint assays enabled sensitive detection of residual disease beyond the resolution of conventional bone marrow diagnostics, providing a refined assessment of treatment response. These findings demonstrate the clinical applicability of patient-specific *MYCN* amplicon breakpoint assays for highly sensitive longitudinal disease monitoring and highlight their potential for dynamic molecular risk assessment throughout therapy.

**Figure 3.**
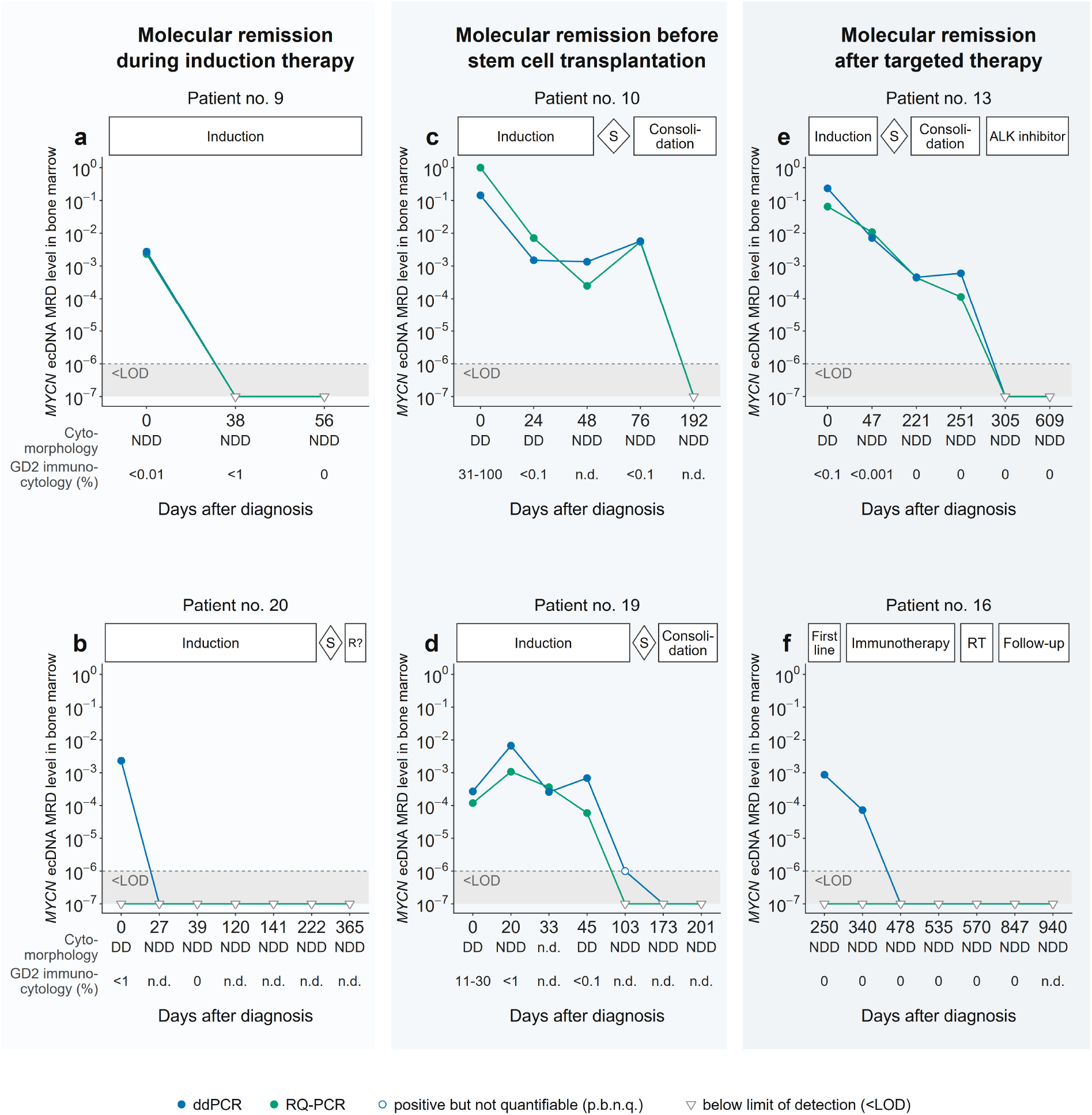
Longitudinal MRD monitoring in serial bone marrow aspirates from patients with high-risk neuroblastoma. Patient-specific *MYCN* amplicon breakpoints quantified by ddPCR and RQ-PCR in mononuclear cells from aspirates collected at diagnosis and throughout therapy. Treatment phases are given above each panel, and cytomorphology and GD2 immunocytology of the same samples beneath it. The dashed line and shaded band mark 10⁻⁶, the lowest tumor DNA fraction reached by the most sensitive assays. Time axes are patient-specific and not to scale. (a, b) Molecular remission during induction therapy. (c, d) Molecular remission before stem cell transplantation. (e, f) Molecular remission after targeted therapy. DD, disseminated disease. NDD, no disseminated disease. n.d., not determined. R?, clinical suspicion of relapse. RT, radiotherapy. S, surgery.

## Discussion

Tumor-informed MRD analysis is not yet standard of care in neuroblastoma. *MYCN* amplification on ecDNA is a defining genomic feature of a large subset of high-risk neuroblastomas. We demonstrate that breakpoints within cancer-specific amplicons, including those on ecDNA, represent uniquely tumor-specific features that can serve as MRD markers complementing established approaches.

The assay developed here for MRD detection in patients with high-risk neuroblastoma is based on patient-specific *MYCN* amplicon breakpoints and combines high-coverage hybrid capture-based panel sequencing for precise breakpoint discovery, with Circle-seq and Nanopore sequencing to resolve amplicon structure, with RQ-PCR and ddPCR for highly sensitive breakpoint detection. Here, we demonstrated MRD monitoring in bone marrow samples. We anticipate that the same breakpoint-specific assays are applicable to cfDNA from blood plasma and other body fluids.

Previous approaches for MRD detection in neuroblastoma used genomic breakpoints (40,41), but were limited by sequencing resolution. Our approach employs targeted panel sequencing to enable high-resolution of breakpoint discovery. In contrast to earlier proof-of-concept studies, we here used clinically applicable workflows (8,23,42). Our approach offers a clinically applicable, integrated solution spanning primary tumor biopsy to a validated MRD assay. Assay setup is required only once per patient and takes approximately 14 working days, from breakpoint discovery to a validated RQ-PCR and ddPCR assay (Figure 4). Every subsequent MRD measurement in that patient then relies on the established assay and can be completed within two working days (Figure 4), making serial monitoring at clinically relevant time points feasible. Thus, our framework for personalized *MYCN* ecDNA MRD assay development provides a foundation for prospective clinical validation and future integration into diagnostic workflows.

**Figure 4.**
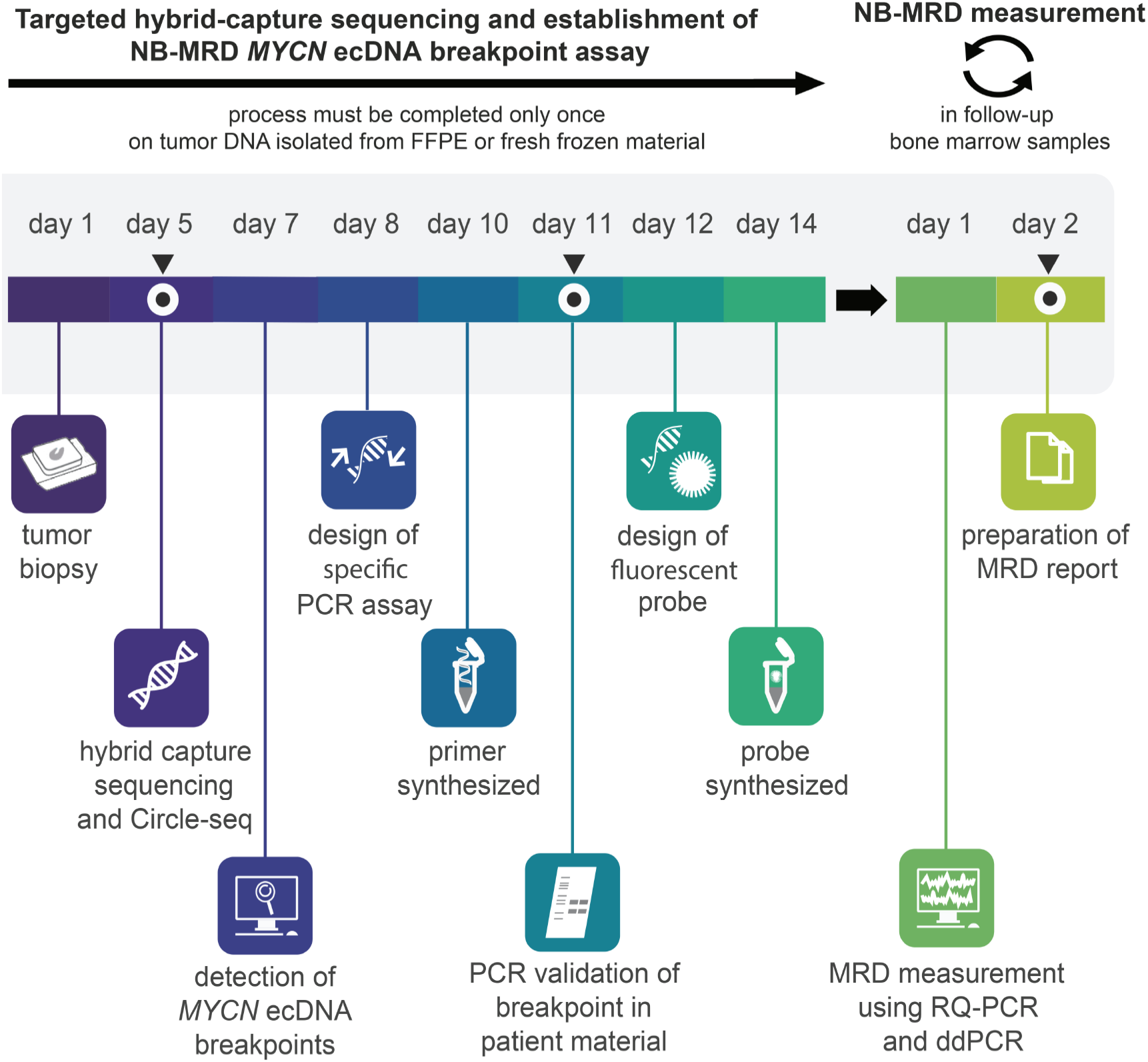
Workflow for clinical implementation of *MYCN* breakpoint-based MRD detection. Breakpoints are mapped from the diagnostic tumor biopsy by hybrid capture-based panel sequencing. Primers and a fluorescent probe are then designed, the breakpoint is validated by PCR and Sanger sequencing, and assay sensitivity is determined in tumor DNA diluted into normal donor DNA. Setup is required once per patient and takes approximately 14 working days. Each subsequent MRD measurement takes two additional days.

A prerequisite for clinical MRD application is marker stability. However, ecDNA exhibits dynamic behavior, including random segregation during mitosis and copy-number heterogeneity (21,43,44), and evolve structurally (22,44,45) as well as numerically (45,46) over the course of treatment. These properties raise the possibility that individual breakpoints could be lost or restructured during or after therapy, or that copy number variance may affect accurate disease quantification. Nevertheless, driver-associated ecDNAs such as those carrying *MYCN* are under strong positive selection (43), are actively retained on mitotic chromosomes (47) and are co-inherited into daughter cells (48), and are therefore expected to persist clonally even as the surrounding amplicon architecture evolves (49). This supports their use as MRD markers while highlighting the importance of longitudinal monitoring. Consistent with this, *MYCN* breakpoints detected at diagnosis were still present at relapse in our cohort, although systematic assessment of amplicon evolution under therapy will be required. Although our current approach focuses on *MYCN*, the same principle applies to other ecDNA-amplified oncogenes, and personalized ecDNA breakpoint detection could therefore be extended beyond neuroblastoma.

PCR-based MRD detection is well established in hematological malignancies, where recurrent fusion genes and clonal immune-receptor rearrangements serve as tumor-specific targets (50,51). Extending this concept to neuroblastoma, we achieved sensitivities of up to 10⁻⁶, exceeding the 10⁻⁴ to 10⁻⁵ typically reached by established MRD assays (52,53). This likely reflects the high copy number of *MYCN* amplicons, which lowers detection thresholds to the point where sensitivity is limited by the number of cells obtainable from a bone marrow aspirate rather than by the assays themselves.

Our findings align with the broader development of tumor-informed MRD strategies in solid tumors, particularly tumor-informed circulating tumor DNA (ctDNA)-based approaches that enable patient-specific disease monitoring and early relapse detection (54,55), including in neuroblastoma, where serial cell-free DNA (cfDNA) analysis detects relapse and actionable targets (56). However, such approaches depend on sufficient tumor DNA shedding, which varies between patients and disease compartments. In this context, our assay provides a complementary strategy that directly interrogates tumor-derived DNA at a primary disease site, enabling highly sensitive detection of residual disease.

Bone marrow involvement is central to neuroblastoma staging and response assessment. Current diagnostic methods, including cytomorphology and GD2 immunocytology (7,57), lack sufficient robustness for reliable MRD detection. Alternative molecular approaches, such as RT-qPCR-based detection of tumor-associated transcripts (15), have shown promise but remain limited by standardization challenges, marker heterogeneity and sample variability. Liquid biopsy approaches have demonstrated feasibility for detecting *MYCN* amplification and tracking clonal evolution in neuroblastoma, both in blood plasma (58,59) and in bone marrow plasma, where cell-free DNA is recoverable at concentrations exceeding those in blood (60), though all such approaches remain dependent on tumor DNA shedding. Our assay differs in the analyte rather than the compartment: it interrogates genomic DNA from cells collected in the bone marrow aspirate, targeting tumor-specific breakpoints present at high copy number and thereby avoiding dependence on shedding. Consistent with previous reports (61), we found that ddPCR provides superior sensitivity and easier assay optimization compared to RQ-PCR, supporting its use as the preferred platform, while RQ-PCR remains a viable alternative.

The bone marrow niche is considered a primary reservoir for residual neuroblastoma cells, supporting its use as the focus for MRD assessment in this study. Breakpoints could also be identified without a targeted panel: WGS (41), Circle-seq (20) and long-read sequencing all resolve amplicon junctions, albeit at higher cost and input requirement and generally without the sequencing depth that targeted capture can provide. Computational reconstruction of focal amplifications has advanced in parallel, both from short-read data (AmpliconArchitect 62; AmpliconClassifier 44) and from long-read data (Decoil 63, CoRAL 64), while Hi-C–based approaches additionally resolve the three-dimensional architecture and regulatory context of amplicons (19,65). Breakpoint-based MRD monitoring has already been demonstrated in ctDNA in neuroblastoma using *TERT* rearrangement breakpoints (66), and cell-free DNA is recoverable from blood, bone marrow, cerebrospinal fluid and urine in these patients (60), so *MYCN* breakpoint-spanning assays could likewise be extended to these compartments. Our approach is therefore intended to complement, rather than replace, established MRD methodologies, and future studies will be required to define their relative performance and optimal integration.

The clinical relevance of MRD detection in neuroblastoma and other solid tumors will need to be established in prospective trials. Based on experience in leukemia, MRD could inform therapy stratification, including decisions on treatment intensity, stem cell transplantation, immunotherapy, and maintenance therapy. MRD levels may also be applied as inclusion criteria or endpoints in future trials testing new agents, enabling more precise measurement of therapy effectiveness.

In summary, we present a personalized MRD detection strategy based on *MYCN* ecDNA breakpoint analysis that integrates advances in ecDNA biology and tumor-informed molecular diagnostics. The assay is highly sensitive, specific, and provides a framework compatible with future clinical implementation following prospective validation. Beyond neuroblastoma, this approach may be applicable to other malignancies characterized by recurrent ecDNAs, supporting broader implementation of ecDNA-based MRD detection strategies in oncology.

## Materials and Methods

### Patients and biomaterial samples and data

Patients had been enrolled in the German neuroblastoma trial (NB2004) or the German Neuroblastoma Registry (NB Registry 2016; 39,67). Written informed consent from patients or parents/guardians was obtained within the trial and registry documentation. Tumor samples were obtained from the German Neuroblastoma Biobank (Cologne) for initial biopsies (8 patients) and biopsies after relapse (3 of these patients). Tumor tissue and bone marrow aspirates were obtained directly from 14 patients (2 with initial and relapse biopsies) treated at Charité Department for Pediatric Oncology and Hematology. Written informed consent was obtained from patients or parents/guardians within the ERA-NET LIQUIDHOPE consortium (TRS-2018-00000715), which obtained ethics approval from the local Charité Ethics Committee (EA2/055/17). The experiments conformed to the principles set out in the WMA Declaration of Helsinki and the Department of Health and Human Services Belmont Report. Serial dilutions were prepared from DNA of tumor samples harboring the breakpoint of interest and from mononuclear cells lacking it, serving as positive and negative spike-in control DNA, respectively, to evaluate MRD assay sensitivity. Control DNA obtained from mononuclear cells from peripheral neuroblastoma-free blood pooled from at least ten healthy individuals was available from the MRD molecular genetics laboratory of the ALL-REZ BFM trials (chair: Arend von Stackelberg, Charité Department for Pediatric Oncology and Hematology, 68). Whole-genome sequencing data in samples from 14 primary *MYCN*-amplified neuroblastomas obtained within our previous studies was used for breakpoint analysis (20,27,28).

### Mononuclear cell purification from bone marrow aspirates

Bone marrow aspirates were collected directly into EDTA-anticoagulated tubes (BD Vacutainer® blood collection tubes). Samples were processed by ficoll density gradient centrifugation as previously published by Krentz/Eckert et al. within 1-4h of collection (69). Aspirates were diluted 1:3 in RPMI 1640 media, and carefully overlaid onto 4ml Biocoll® (Biochrom) per 5ml diluted aspirate, then separated by centrifugation at 2400rpm for 20min at 20°C and decelerating without brake. The bone marrow mononuclear cell fraction (BM-MNCs) was aspirated out of the gradient and washed twice with equal volumes of RPMI 1640 followed by sedimentation at 1500rpm for 10min at 20°C. If samples were contaminated with erythrocytes, they were lysed with ammonium chloride, before washing mononuclear cells again twice with 1x PBS and centrifugations at 1500rpm for 10min at 20°C. Mononuclear cells were quantified, and DNA was isolated from 1 x 10^6^ to 4 x 10^6^ cell aliquots using the NucleoSpin Tissue kit (Macherey-Nagel GmbH & Co. KG) according to the manufacturer’s directions before storing at - 20°C until use in hybrid capture-based panel sequencing, droplet digital or real-time quantitative PCR.

### Cell Lines

Neuroblastoma cell lines KELLY, LAN-5, NGP, TR-14, SK-N-DZ, IMR-5, SK-N-BE and CHP-212 (all *MYCN*-amplified) were analysed in this study. CHP-134 (*MYCN*-amplified) and HeLa cervix adenocarcinoma were used as positive and negative PCR controls, respectively, and were not included in the breakpoint analyses. The neuroblastoma cell lines KELLY and LAN-5 were obtained from the German Collection of Microorganisms and Cell Cultures GmbH (DSMZ; Braunschweig, Germany). NGP was kindly provided by F. Speleman (Cancer Research Institute Ghent, Ghent, Belgium). TR-14 was kindly provided by J. J. Molenaar (Princess Máxima Center for Pediatric Oncology, Utrecht, Netherlands), SK-N-DZ was kindly provided by A. Künkele and SK-N-BE and CHP-212 was kindly provided by J.H. Schulte (both then at Charité - Universitätsmedizin Berlin), and IMR-5, CHP-134 and HeLa cervix adenocarcinoma were kindly provided by A. Schramm (Medizinische Fakultät, Universitätsklinikum Essen, Essen, Germany). Cells cultured in RPMI 1640 or DMEM (Thermo Fischer Scientific, Waltham, MA, USA) supplemented with 10% fetal calf serum (GE Healthcare, Chicago, IL, USA), 1% penicillin (100 U/ml, Thermo Fischer Scientific) and 1% streptomycin (100 μg/ml, Thermo Fischer Scientific) at 37°C and 5% CO_2_. Master stocks for all cell lines were authenticated by short tandem repeat DNA typing by Idexx Bioresearch (Westbrook, ME, USA) or Multiplexion (Heidelberg, Germany) before short-term cell culture and subsequent DNA extraction. Cultures were routinely tested for mycoplasma using the MycoAlert Mycoplasma Detection Kit (Basel, Switzerland).

### DNA fluorescence in situ hybridization

Fresh-frozen neuroblastoma samples were cut into sections (6µm) using a cryostat microtome (SLE Medical GmbH). Tumor cell content was determined on hematoxylin and eosin-stained sections by a pathologist. Neuroblastoma cell lines were seeded separately in a four chamber Nunc™ Lab-Tek™ II Chamber Slide™ system (Sigma-Aldrich) and cultured for at least 48h. Afterwards, the media was discarded and the slides were washed once with 1x PBS and fixed in a 4% Paraformaldehyde in 1xPBS (neoLab Migge GmbH) for 10min. Slides were rinsed in 1× PBS and air-dried. DNA FISH analysis was performed according to neuroblastoma trial protocol standards(70). To detect a *MYCN* gene amplification, a DNA probe complementary to chr.2p24 (Vysis LSI N-MYC SG/CEP2 SO, start at chr2:15,957,690, stop at chr2:16,163,618; Vysis, Abbott Laboratories) was used. A CEP2 probe (Vysis, Abbott Laboratories) which binds a repetitive sequence in the centromeric region was added as a reference. FISH was performed in a dual-color procedure following the manufacturers’ instructions. Sections were covered using ProLong™ Gold Antifade Mountant (Thermo Fischer Scientific), including nuclear counterstaining with DAPI (4′-6-diamidino-2-phenylindole) blue-fluorescent dsDNA binding dye. Fluorescence images were captured on a Leica DMRA microscope (Leica Camera) equipped with JVC KY F-75 digital camera (JVC) using Diskus Viewer software (Hilgers Technisches Buero e.K.). Results were analyzed according to recommendations from the European Neuroblastoma Quality Assessment group(71).

### Re-analysis of next-generation tumor sequencing data for detection of *MYCN* structural variants

Publicly available whole-genome sequencing data (20,27,28) were reanalyzed for structural variants in proximity to *MYCN*, using novoBreak software (v1.1) with default parameter settings. Variant calls were stringently filtered to include only breakpoints which had strong support in the tumor samples and were absent from matched control samples. Breakpoints were required to have a quality score >50, at least 3 high-quality spanning reads from the tumor sample (<1 such read in the matched control) and at least 10 high-quality discordant read pairs in the tumor (<3 in the matched control).

### Tumor DNA extraction for hybrid capture-based panel sequencing

Fresh-frozen or formalin-fixed, paraffin-embedded tumor material was histologically evaluated by an experienced pathologist, and only regions with at least 10% tumor cell content were macrodissected. Tumor cells were enriched by laser microdissection (Leica LMD 6500, Leica Microsystems, Wetzlar, Germany) for cases with limited tumor cell content. DNA was automatically extracted using a Maxwell Instrument (Promega, Fitchburg, WI, United States), then sheared mechanically by ultrasonic acoustic energy (Covaris ultrasonicator, Covaris, Matthews, NC, USA) before subjecting 100-200ng to our neuroblastoma hybrid capture sequencing panel(31).

### Neuroblastoma hybrid capture-based panel sequencing

The NucleoSpin Tissue kit (Macherey-Nagel GmbH & Co. KG) was used to isolate DNA from neuroblastoma cell lines for hybrid capture-based panel sequencing. Tumor DNA was isolated from macrodissected cells from 10-15 10μm sections of formalin-fixed, paraffin-embedded or fresh frozen tumor material as described in the Supplemental Methods. From each sample, 100-200ng were subjected to our custom neuroblastoma hybrid capture sequencing panel (NEO New Oncology GmbH, Cologne, Germany) for neuroblastoma to detect point mutations, small insertions and deletions, copy-number alterations and rearrangements/gene fusions and to map the *MYCN* amplicon in a single assay(31). Adapters were ligated to sheared sample DNA, and individual genomic regions of interest were enriched using complementary bait sequences (hybrid capture procedure). Selected baits ensured optimal coverage of all relevant genomic regions. After enrichment, targeted fragments were amplified (clonal amplification) and sequenced in parallel with an average mean sequencing depth of 1000-2500x (unfiltered reads). Sequencing data were processed using the NEO New Oncology custom analysis pipeline. Briefly, the quality of the raw sequencing reads was assessed, and the reads were aligned to the GRCh37 (hg19) human reference genome assembly. Discordantly aligned read pairs were subsequently extracted, and structural variants and their breakpoints were localized. Breakpoint position and supporting data (spanning and encompassing sequencing reads) were recorded for all detected alterations.

### Circular DNA isolation, purification and sequencing

Circular DNA isolation and purification was performed similarly to previous reports of Circle-seq (20,72). DNA content was measured with a NanoDrop 2000 Spectrophotometer (Thermo Fisher Scientific) and a Qubit 3.0 Fluorometer (Thermo Fisher Scientific). Amplified circular DNA was sheared to an average fragment size of 150–200bp using an S220 focused ultrasonicator (Covaris). Libraries for next-generation sequencing were prepared using the NEBNext Ultra DNA Library Kit for Illumina according to the manufacturer’s protocol (New England Biolabs). Libraries were sequenced on MiSeq instruments with 2 × 150 bp paired-end reads, HiSeq 4000 instruments with 2 × 125 bp paired-end reads or NextSeq 500 instruments with 2 × 150 bp paired-end reads (all Illumina).

### Circle-seq analysis

Circle-seq analysis was done in accordance with Koche et al. 2020 (20). Reads were 3′ trimmed for both quality and adapter sequences, with reads removed if the length was less than 20 nucleotides. Reads were aligned to the human hg19 reference assembly using the Burrows–Wheeler Aligner MEM v.0.7.15 (default parameters). PCR and optical duplicates were removed with Picard v.2.16.0. The aligned BAM files were analyzed in two ways. All read pairs and split reads containing any outward-facing read orientation, indicating potential circles, were placed in a new BAM file. Genomic segments enriched for signal over background were detected in the ‘all reads’ BAM file using variable-width windows from Homer v.4.11 findPeaks (http://homer.ucsd.edu/), and the edges of these enriched regions were intersected with the ‘circle only’ BAM file to quantify the number of circle-supporting reads. To determine the thresholds for significance of real circles versus background noise, matched WGS data were used to determine the background distribution of circle-oriented reads in regions without circles that were matched for length and nucleotide composition. An empirical P value of 0.01 was used to filter putative circles, and regions passing this filter were then used for downstream analysis.

### Nanopore DNA Sequencing

DNA was extracted using the MagAttract HMW DNA Kit (Qiagen N.V., Venlo, Netherlands). The Ligation Sequencing Kit (SQK-LSK109, Oxford Nanopore Technologies Ltd., Oxford, UK) was used for library preparation(19). Nanopore sequencing was performed on a R9.4.1 MinION flowcell (FLO-MIN106, Oxford Nanopore Technologies Ltd., Oxford, UK). We used Guppy 2.3.7 (Oxford Nanopore Technologies Ltd., Oxford, UK) with default parameters for basecalling, NGMLR 0.2.7 (73) for read alignment and Sniffles 1.0.11 (73) to call structural variants. Coverage was computed as unnormalized number of reads in 20bp bins using deepTools 3.3.0 (74).

### Design of PCR assays along *MYCN* amplicon breakpoints

The exact sequence of about 200 base pairs surrounding the breakpoint up and downstream was exported using IGV (Integrative Genomics Viewer). A forward primer was designed upstream of the breakpoint, and a reverse primer was designed downstream to the breakpoint for the opposite strand. PCR product size was at least 70 nucleotides and up to 350 nucleotides in length. We designed at least 3 primer pairs for each breakpoint for a confirmation of the sequence. The best primer pair was chosen for breakpoint detection using RQ-PCR or ddPCR, and an additional TaqMan probe of approximately 20 nucleotides and using a fluorescein and Black Hole Quencher 1 dye (5’-6-fluorosine amidite, FAM - BHQ-1-3’) was designed. Primer pairs flanked the breakpoint, and the probe was positioned as not to cover the breakpoint. The designed primers and TaqMan probes were applied for the breakpoint-specific real-time quantitative PCR (RQ-PCR) and the breakpoint-specific digital droplet PCR (ddPCR).

### Conventional PCR for breakpoint validation

*MYCN* amplicon breakpoints were validated in DNA prepared as for neuroblastoma hybrid capture sequencing(31) from tumor samples, mononuclear cells or neuroblastoma cell lines using GoTaq Hot Start Polymerase (Promega, Madison, WI, USA) in a standard 30-cycle PCR reaction. PCR products were analyzed by QIAxcel advanced automatic electrophoresis (Qiagen, Hilden, Germany) using a DNA high-resolution cassette with the QX 15bp/3kb alignment marker and QX pUC18/HaeIII DNA size marker according to the manufacturer’s instructions. Each PCR run included a *MYCN* amplification control (DNA from Kelly or CHP-134 *MYCN*-amplified neuroblastoma cell lines) and 2 negative controls (DNA from the HeLa cervical cancer cell line and DNA from mononuclear cells pooled from at least 10 healthy donors 33,68). PCR products were sequenced by Eurofins Genomics GmbH (Ebersberg, Germany) or LGC Genomics GmbH (Berlin, Germany) directly and after subcloning into pUC18.

### Breakpoint-specific real-time quantitative PCR

Real-time quantitative PCR was performed as previously described for MRD detection in patients with acute lymphoblastic leukemia (33,75). DNA was isolated from tumors, mononuclear cells or neuroblastoma cell lines as for neuroblastoma hybrid capture sequencing(31). At least one breakpoint of the patient-specific *MYCN* amplicon was selected for each patient (or cell line used for assay development or validation), and the primer pair and a 5’-6-FAM-BHQ-1-3’ probe were designed for RQ-PCR. PCR-amplified sequences were detected via the FAM tag in real time. Assay sensitivity was determined from serial dilutions of tumor DNA into DNA from healthy donors down to 10⁻⁶ and is expressed as the lowest tumor DNA fraction reproducibly detected. The StepOnePlus real-time PCR System (Thermo Fischer Scientific Scientific, Waltham, MA, USA) was used for RQ-PCR using a mix containing 670 ng DNA in a total volume of 13.3µl, comprising 3.1µl ddH_2_O, 3µl MgCl_2_ 50mM, 2µl 10x Buffer, 2µl 2mM dNTPs, 1µl each 10pmol/µl forward/reverse primer, 0.5µl bovine serum albumin, 0.5µl 10pmol/µl dual-labeled fluorescent hydrolysis probe, (6-carboxyfluorescein, 6-FAM, and the BHQ1 quencher), 0.2µl 5U/µl Platinum Taq DNA Polymerase and running on the program, 94°C for 5min followed by 50 cycles of 94°C for 8sec and 60-65°C (optimized primer annealing temperature) for 23sec, followed by cool-down to 4°C. Data was analyzed using the StepOnePlus Software Version 2.3 (Applied Biosystems, Foster City, CA, USA), adapted to the guidelines of Van der Velden (33) and according to EuroMRD Consortium guidelines(76). The housekeeping gene, hemoglobin subunit beta, was used as a reference gene for normalization as previously described(33). For quantification of *MYCN* amplicon breakpoints, the ratio of cells harboring:lacking the breakpoint in each patient sample was calculated using standard curve equations(33). All assays were conducted using technical triplicates.

### Breakpoint-specific droplet digital PCR

Droplet digital PCR was performed using RQ-PCR primer/probe sets on DNA isolated as for RQ-PCR. Droplet PCR results were quantified in single-channel detection mode (6-FAM-tag) for absolute quantification on a Bio-Rad QX200 Droplet Reader. Each ddPCR reaction contained 670 ng DNA and was analyzed in technical triplicates. Assay sensitivity was determined from serial dilutions of tumor DNA into pooled normal donor DNA and is expressed as the lowest tumor DNA fraction reproducibly detected. Reactions were prepared as follows: Premix was made containing 2.1µl double-distilled H_2_O, 12µl ddPCR probe supermix without dUTP (Bio-Rad Laboratories, Inc., Hercules, CA, USA), 1.7µl each of 10 pmol/µl forward/reverse primer and 0.5µl of 10 pmol/µl FAM-BHQ1-labeled probe. Each reaction consisted of 18µl PCR premix and 2µl of denatured sample DNA (600-670ng sample DNA was denatured at 95°C for 10min). The PCR reaction was partitioned into droplets in 70µl Droplet Generation Oil (Bio-Rad) using the QX200 Droplet Generator (Bio-Rad), and 40µl manually transferred to 1 well in a 96-well PCR plate in a T100 Thermal Cycler running the program: 95°C for 10min followed by 40 cycles of 94°C for 30sec and 60-70°C (optimized primer annealing temperature) for 1min, then ramped up to 98°C (2°C/sec) for 10min before cool-down to 4°C. The housekeeping gene, hemoglobin subunit beta, was used as a reference gene for normalization among samples. Data was analyzed using QuantaSoft 1.7.4.0917 or QuantaSoft™ Analysis Pro 1.0.596 software (Bio-Rad) with MRD level assessment as previously published for acute lymphoblastic leukemia and in agreement with current ddPCR recommendations of the EuroMRD Consortium(77).

### Statistical analysis

Analyses were performed in GraphPad Prism v.7. Correlation between absolute *MYCN* copy number and the number of detected breakpoints was assessed by Spearman rank correlation, separately for cell lines and primary tumors. *MYCN* copy number in samples with ecDNA versus HSR-integrated amplicons was compared using the Mann–Whitney U test. Assay sensitivities, expressed as log₁₀ dilution steps, obtained by RQ-PCR and ddPCR were compared across the same breakpoints using the Wilcoxon signed-rank test for paired data. All tests were two-sided; p < 0.05 was considered significant. Given the descriptive nature of these comparisons, no adjustment for multiple testing was applied.

### CRediT authorship contribution statement

Annabell Szymansky: Conceptualization, Methodology, Investigation, Formal analysis, Software, Data curation, Visualization, Writing – original draft, Writing – review and editing. Falk Hertwig: Conceptualization, Investigation, Writing – review and editing. Annika Winkler: Investigation, Writing – review and editing. Rocío Chamorro González: Formal analysis, Software. Konstantin Helmsauer: Formal analysis, Software. Anna-Marie Schroeer: Investigation. Melanie J. Witthauer: Investigation. Eliz Karaman: Investigation. Marco Lodrini: Conceptualization, Resources, Visualization. Jasmin Wünschel: Investigation, Resources. Nadine Wittstruck: Investigation, Resources. Steffen Fuchs: Resources. Jessica Theissen: Resources. Barbara Hero: Resources. Richard Koche: Methodology. Malwine J. Barz: Investigation, Visualization. Lukas C. Heukamp: Investigation, Methodology, Funding acquisition. Kathy Astrahantseff: Writing – review and editing. Karin Schmelz: Resources. Patrick Hundsdoerfer: Resources. Joern Toedling: Formal analysis, Software, Writing – review and editing. Angelika Eggert: Funding acquisition, Supervision, Project administration. Cornelia Eckert: Conceptualization, Methodology, Supervision, Project administration, Writing – original draft. Bastiaan Spanjaard: Visualization, Writing – review and editing. Hedwig E. Deubzer: Supervision, Resources, Funding acquisition, Project administration, Writing – review and editing. Matthias Fischer: Supervision, Resources, Funding acquisition. Anton G. Henssen: Conceptualization, Supervision, Funding acquisition, Writing – original draft, Writing – review and editing. Johannes H. Schulte: Conceptualization, Resources, Supervision, Funding acquisition, Project administration, Writing – original draft.

All authors read and approved the final manuscript.

### Data availability

Targeted sequencing data from tumors are deposited in the European Genome-phenome Archive (http://ega-archive.org/studies/) under accession numbers EGAS50000001581 (study ID: 2476), EGAS00001007365 (dataset ID: EGAD00001011088) and EGAS00001004275. Targeted sequencing data from neuroblastoma cell lines are in the NCBI Sequence Read Archive (http://www.ncbi.nlm.nih.gov/Traces/sra) under BioProject numbers PRJNA1301554 (submission ID: SUB15515512) and PRJNA979802. Nanopore sequencing data of the *MYCN* amplicon were reported previously (19) and are available in the NCBI Sequence Read Archive (BioProject: PRJNA622577). Publicly available whole-genome sequencing data reanalyzed to define the *MYCN* breakpoint target region are deposited in the European Genome-phenome Archive under accession numbers EGAS00001001308, EGAS00001003244 and EGAS00001004022. Original RQ-PCR and ddPCR data are available upon request.

## Acknowledgements

The authors thank the patients and their parents, who agreed to take part in this study, all physicians involved in biosample collection and our hematological laboratory technicians, Constanze Passenheim, Nadine Sachs and Simone Roßa for their excellent work. We thank Jutta Proba, Anke Schaubitzer, Iris Gruska and Daniel Hirsch for technical support. We thank the German Neuroblastom Biobank for providing tumor samples and the Institut für Hämatopathologie Hamburg for targeted sequencing. We thank Elias Rodriguez-Fos for helpful discussion of the data.

