## Supplemental Figures for "Quantitative and sensitive neuroblastoma minimal residual disease detection using extrachromosomal DNA (ecDNA) breakpoints": SuppFigure3_MYCN-ini-rez.pdf

Patient 2 - Breakpoint no. 7

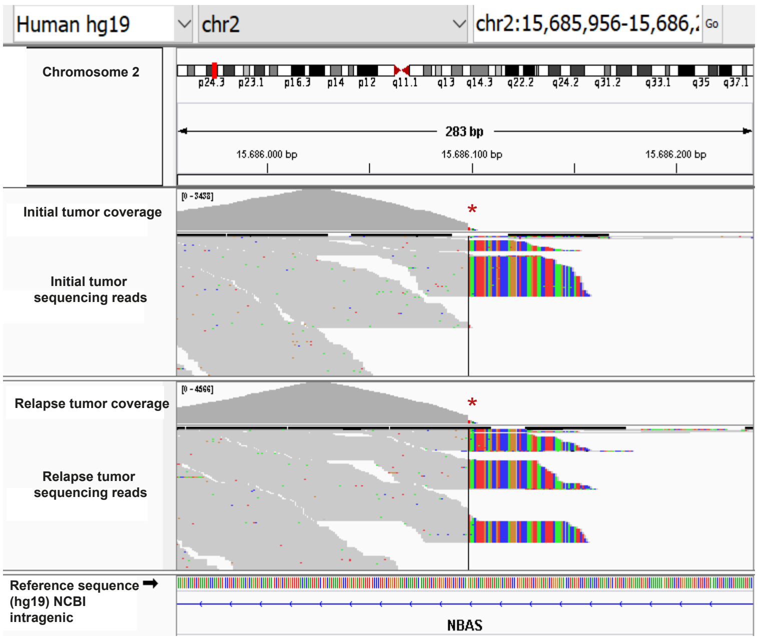


Patient 5 - Breakpoint no. 13

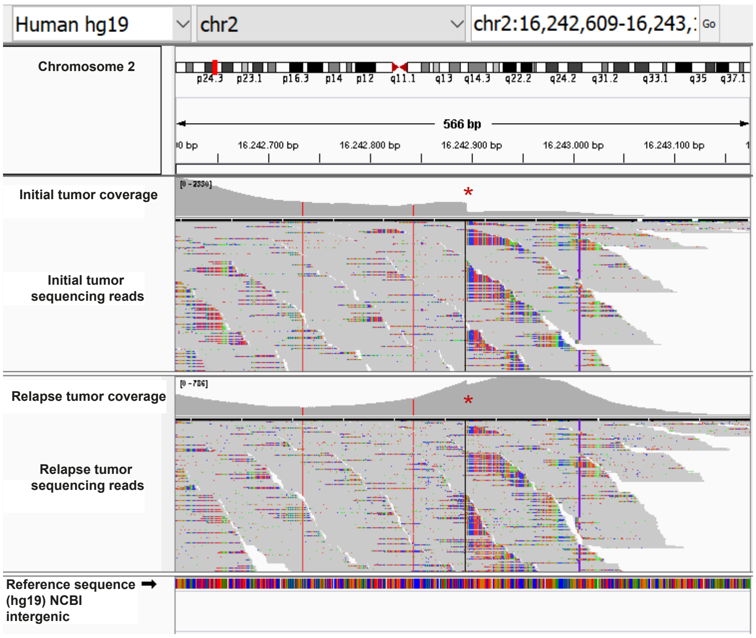


Patient 8 - Breakpoint no. 19

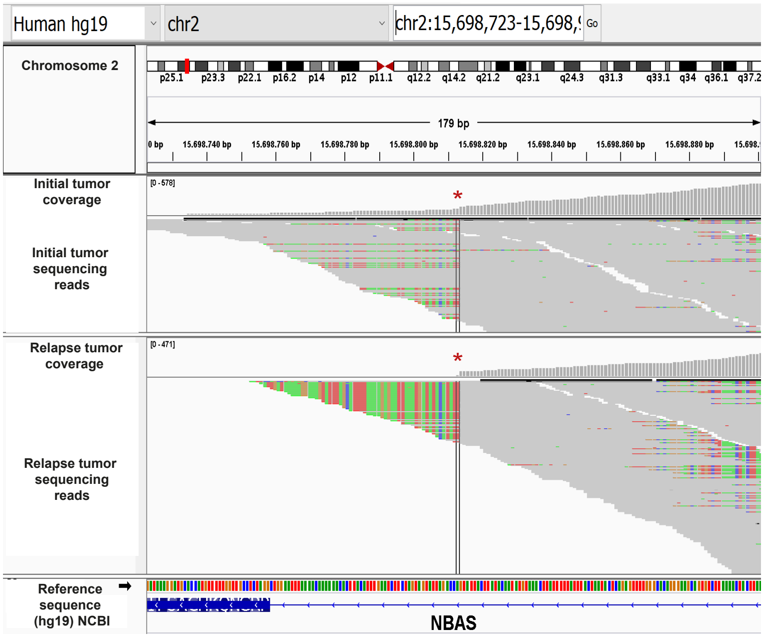


Patient 15 - Breakpoint no. 39

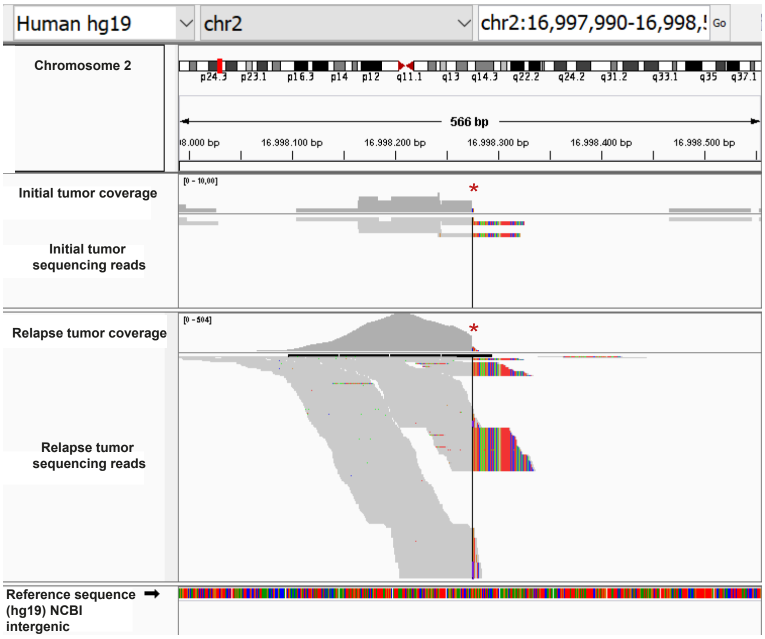


Patient 16 - Breakpoint no. 43

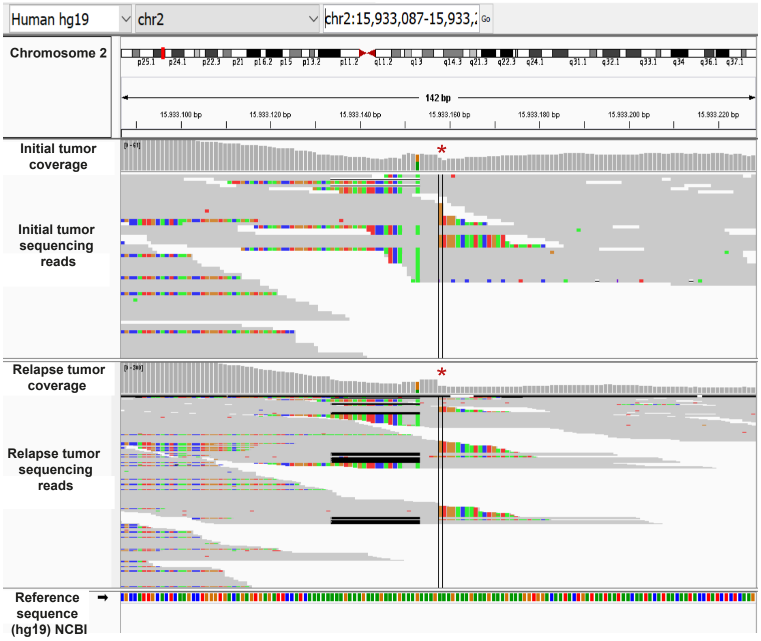
