## Supplemental Figures for "Quantitative and sensitive neuroblastoma minimal residual disease detection using extrachromosomal DNA (ecDNA) breakpoints": SuppFigure4_TR14_circleBPs.pdf

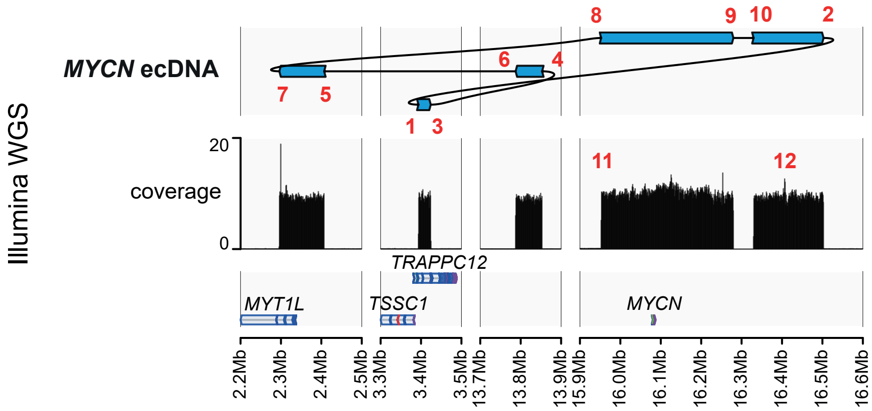


| Breakpoint number | Chromosome start | Breakpoint Start | Orientation start | Breakpoint number | Chromosome end | Breakpoint end | Orientation end |
|---|---|---|---|---|---|---|---|
| 1 | chr2 | 3393721 | - | 2 | chr2 | 16502814 | + |
| 3 | chr2 | 3424107 | + | 4 | chr2 | 13852991 | + |
| 5 | chr2 | 2406951 | + | 6 | chr2 | 13787343 | - |
| 7 | chr2 | 2295975 | - | 8 | chr2 | 15952491 | - |
| 9 | chr2 | 16279724 | + | 10 | chr2 | 16329724 | - |
| 11 | chr2 | 15966854 | + | 12 | chr2 | 16400242 | - |

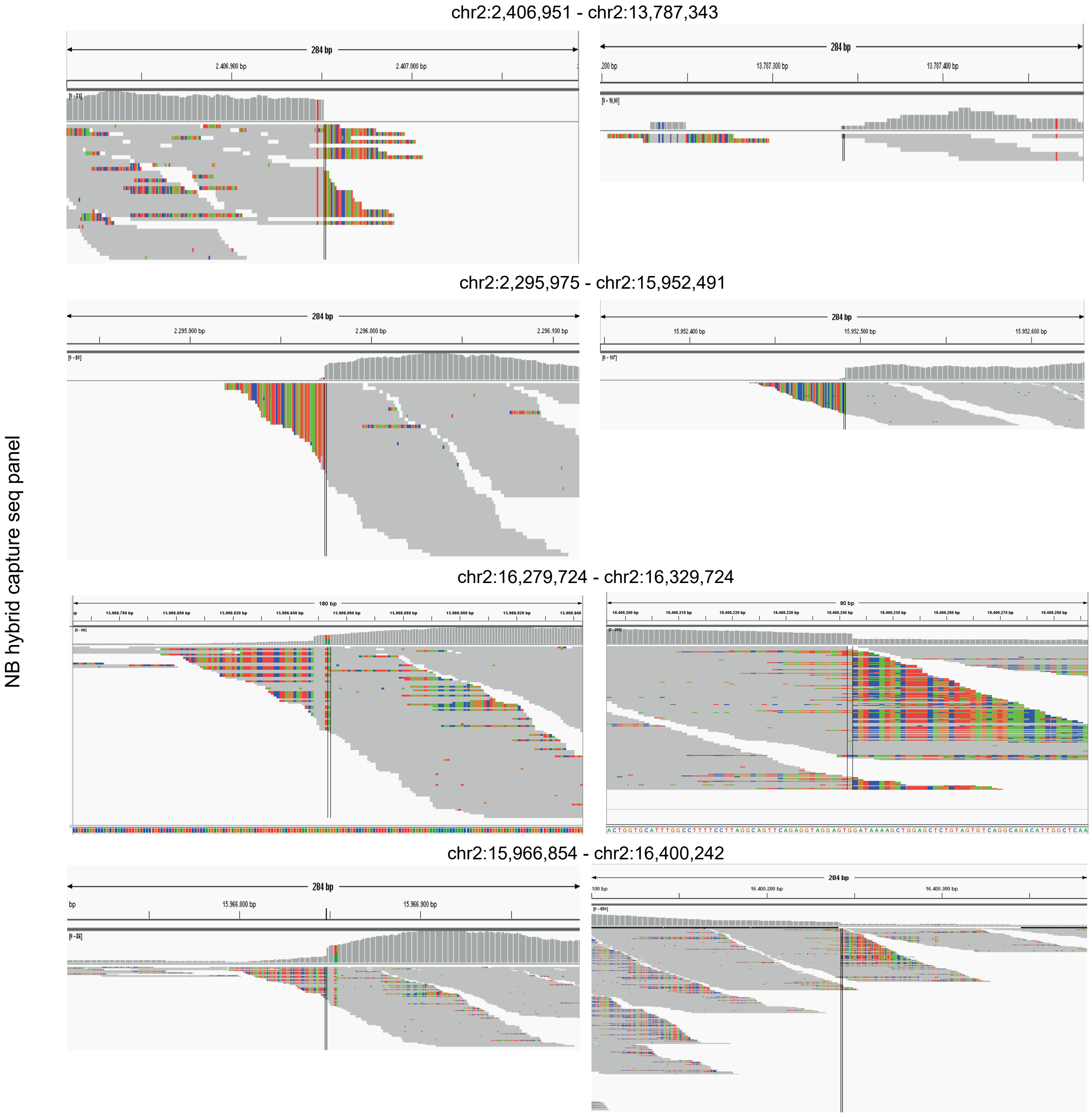
