## Supplemental Figures for "Quantitative and sensitive neuroblastoma minimal residual disease detection using extrachromosomal DNA (ecDNA) breakpoints": SuppFigure6_sangerpatient_continued.pdf

patient 12

chr2:16084511 chr2:16697303

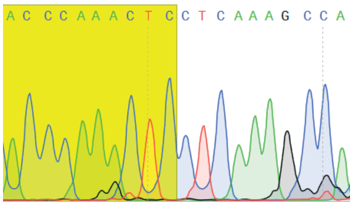

TGCATTTCCCACCCAAACTCC
TCAAAGCCACCAATAATAA

patient 12

chr2:16082975 chr2:16180006

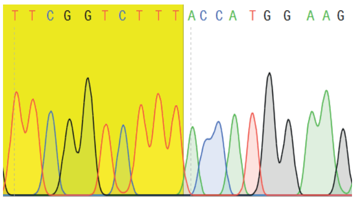

CGGACCCGAGTTCGGTCTTTA
CCATGGAAGATGAACATTT

patient 12

chr2:16105859 chr2:16220058

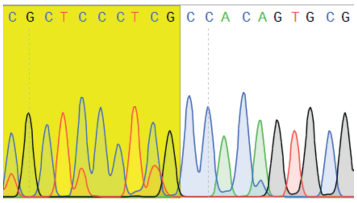

AGTGTCTCATCGCTCCCTCGC
CACAGTGCGGAGGGTTGAG

patient 13

chr2:6472899 chr2:16077648

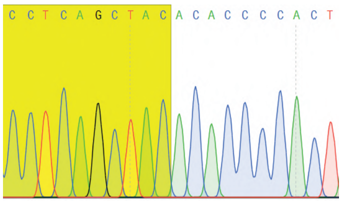

CAGTCAGGTCCCTCAGCTACA
CACCCCACTAGCATCATGA

patient 13

chr2:15485584 chr2:15829866

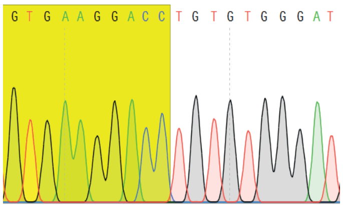

AACAAGGGATGTGAAGGACCT
GTGTGGGATGTGTGTTCT

patient 13

chr2:15089860 chr2:16574270

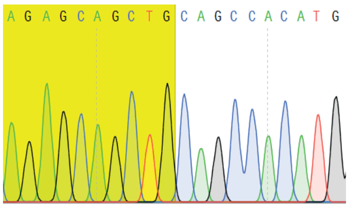

CAGCCAGGACAGAGCAGCTGC
AGCCACATGGATGGAACTG

patient 13

chr2:16895922 chr2:16896040

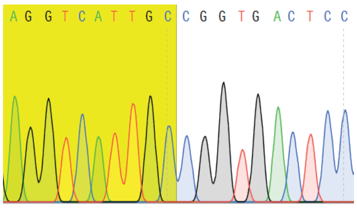

TGAGTGGGTGAGGTCATTGCC
GGTGACTCCCTGCTGCCAC

patient 14

chr2:15276043 chr2:15999479

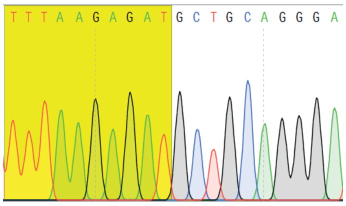

CTCATAATATTTTAAGAGATG
CTGCAGGGATGTCTTATTCT

patient 14

chr2:15857004 chr2:16426962

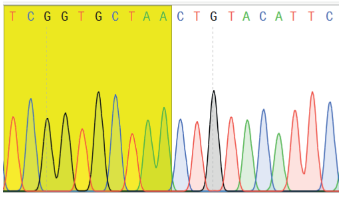

TATGACTAACTCGGTGCTAAC
TGTACATTCATTCACTCAT

patient 14

chr2:16077433 chr2:16472680

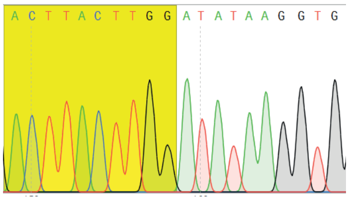

GGATACCTTGACTTACTTGGA
TATAAGGTGGCTGCAGAAT

patient 14

chr2:16129580 chr2:16717055

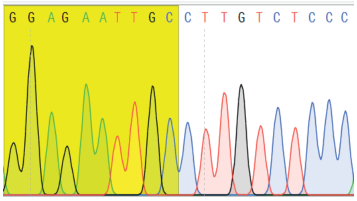

GGCTGAGGCAGGAGAATTGC
CTTGTCTCCCAAAGTGCTGG

patient 14

chr2:15392011 chr2:15814812

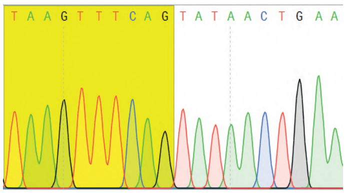

TGCAGCCCAGTAAGTTTCAGT
ATAACTGAAACTATAGAGG

patient 15

chr2:16663369 chr2:17253962

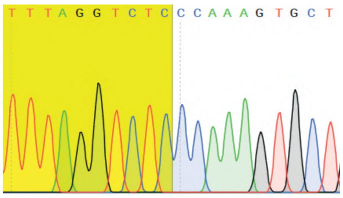

GAGTATTCATTTTAGGTCTCC
CAAAGTGCTGGGATTACAG

patient 15

chr2:16075579 chr2:17068299

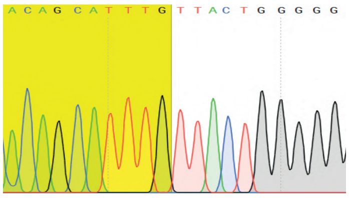

GCTTGTAATCACAGCATTTGT
TACTGGGGGGTTGATGCAG

patient 15

chr2:16395748 chr2:16998274

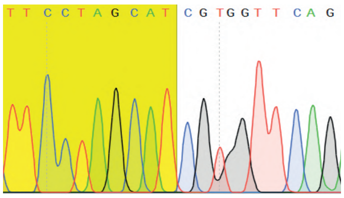

TCTGGAAAGCTTCCTAGCATC
GTGGTTCAGAGATGGCCTCT

patient 16

chr2:15736169 chr2:16171673

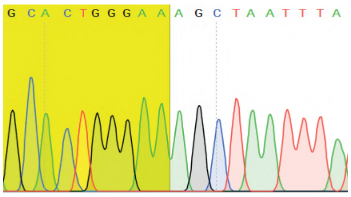

ATTGCTCTTAGCACTGGGAAA
GCTAATTTATTGTATTTTT

patient 16

chr2:15932872 chr2:15933444

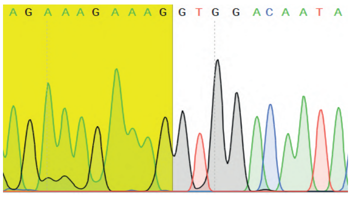

AAAGAAAGAAAGAAAGAAAGG
TGGACAATACTGATAATAA

patient 17

chr2:15858551 chr2:16459069

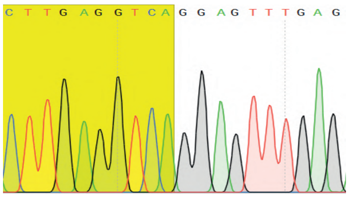

CTCAAACTCCTGACCTCAAGT
GAAATTGCAGAAGCTCTCT
