## Supplemental Figures for "Quantitative and sensitive neuroblastoma minimal residual disease detection using extrachromosomal DNA (ecDNA) breakpoints": Supplemental_Figure-Legends.pdf

**Supplemental Figure S1. *MYCN* amplification on extrachromosomal DNA shown by fluorescence in situ hybridization.** Interphase FISH with a *MYCN* probe and a chromosome 2 centromeric *CEP2* reference probe, with DAPI counterstain, for the three neuroblastoma cell lines and the nine primary tumors in which the signal pattern was dispersed. Each panel is annotated with its own channel colors: *MYCN* green and *CEP2* red throughout, except patient no.5, where *MYCN* is red and *CEP2* green. Arrows mark representative dispersed *MYCN* signals, the pattern consistent with amplification on extrachromosomal DNA. Clustered signals, indicating intrachromosomal amplification as a homogeneously staining region, were observed in three further tumors and in five cell lines and are not shown. Scale bars, 50 µm.

**Supplemental Figure S2. Breakpoint density around *MYCN* and orthogonal validation of the hybrid capture assay.** (a) Frequency of genomic breakpoints within two megabases around *MYCN* identified by whole-genome sequencing of 14 primary *MYCN*-amplified neuroblastomas, in 200 kb bins. Beneath the histogram are the positions of the hybrid capture probes tiling the region, the coordinates on chromosome 2 and the genes it contains. The red box on the ideogram marks the target region. (b) The same region and layout for the breakpoints detected with the hybrid capture sequencing assay, with a smoothed density function overlaid on the histogram. (c) Circle-seq coverage across chr2:10,894,994–17,451,251 in the initial tumor biopsy of patient #9. The arc and the red triangles mark the junction chr2:13,490,249 to chr2:15,815,066. (d) Reads from the neuroblastoma hybrid capture sequencing panel at both sides of the same junction in the same sample, confirming it independently of the Circle-seq data.

**Supplemental Figure S3. *MYCN* amplicon breakpoints in paired initial and relapse tumor samples.** Read-level evidence from hybrid capture sequencing for one breakpoint in each of the five patients with paired biopsies obtained at diagnosis or during therapy and at relapse (patients #2, #5, #8, #15 and #16). For each patient, the coverage track and the aligned reads of the initial tumor are shown above those of the relapse tumor, and the breakpoint is marked by an asterisk. Soft-clipped bases appear as colored mismatches. The panels for patients #2, #5, #8 and #16 show breakpoints detected in both samples. The panel for patient #15 shows breakpoint 39, one of the three breakpoints detected only in the relapse biopsy.

**Supplemental Figure S4. Reconstruction and read-level validation of the *MYCN* amplicon in the cell line TR-14.** Top: reconstruction of the *MYCN* ecDNA in TR-14 with the twelve breakpoint ends numbered, shown above the Illumina whole-genome sequencing coverage and the gene track for the four chromosome 2 segments it comprises. Middle: chromosome, position and orientation of both ends of each of the six junctions. Bottom: aligned reads from the neuroblastoma hybrid capture sequencing panel at both sides of four of these junctions, confirming them independently of the whole-genome sequencing data.

**Supplemental Figure S5. Sanger sequencing validation of breakpoints in neuroblastoma cell lines.** Electropherograms of breakpoint-spanning PCR products confirming 21 breakpoints detected in the 8 neuroblastoma cell lines. Each panel is labeled with the cell line and the two genomic positions joined at the junction, and the sequence spanning the junction is given beneath the trace.

**Supplemental Figure S6. Sanger sequencing validation of breakpoints in primary neuroblastomas.** Electropherograms of breakpoint-spanning PCR products confirming 42 of the 47 breakpoints detected in the 22 primary tumors, for patients #1 to #11 on the first page. Each panel is labeled with the patient number and the two genomic positions joined at the junction, and the sequence spanning the junction is given beneath the trace. The figure is continued on a second page with patients #12 to #17.

**Supplemental Figure S7. Longitudinal MRD monitoring in the remaining patients of the cohort.** Patient-specific *MYCN* amplicon breakpoints quantified by ddPCR (blue) and RQ-PCR (green) in serial bone marrow aspirates from the eight patients not shown in Figure 3, but displayed as in Figure 3. (a) Patient #11. (b) Patient #12. (c) Patient #14. (d) Patient #15. (e) Patient #17. (f) Patient #18. (g) Patient #21. (h) Patient #22. Time axes are patient-specific and not to scale, and give days after diagnosis except in (c), which gives days after relapse, and (d), which gives days after the start of the first N4 cycle. DD, disseminated disease. NDD, no disseminated disease. n.d., not determined. p.b.n.q., positive but not quantifiable. S, surgery. SCT, stem cell transplantation.
