## Supplementary figures and images for "Quantitative and sensitive neuroblastoma minimal residual disease detection using extrachromosomal DNA (ecDNA) breakpoints"

### SuppFigure1_FISH.pdf

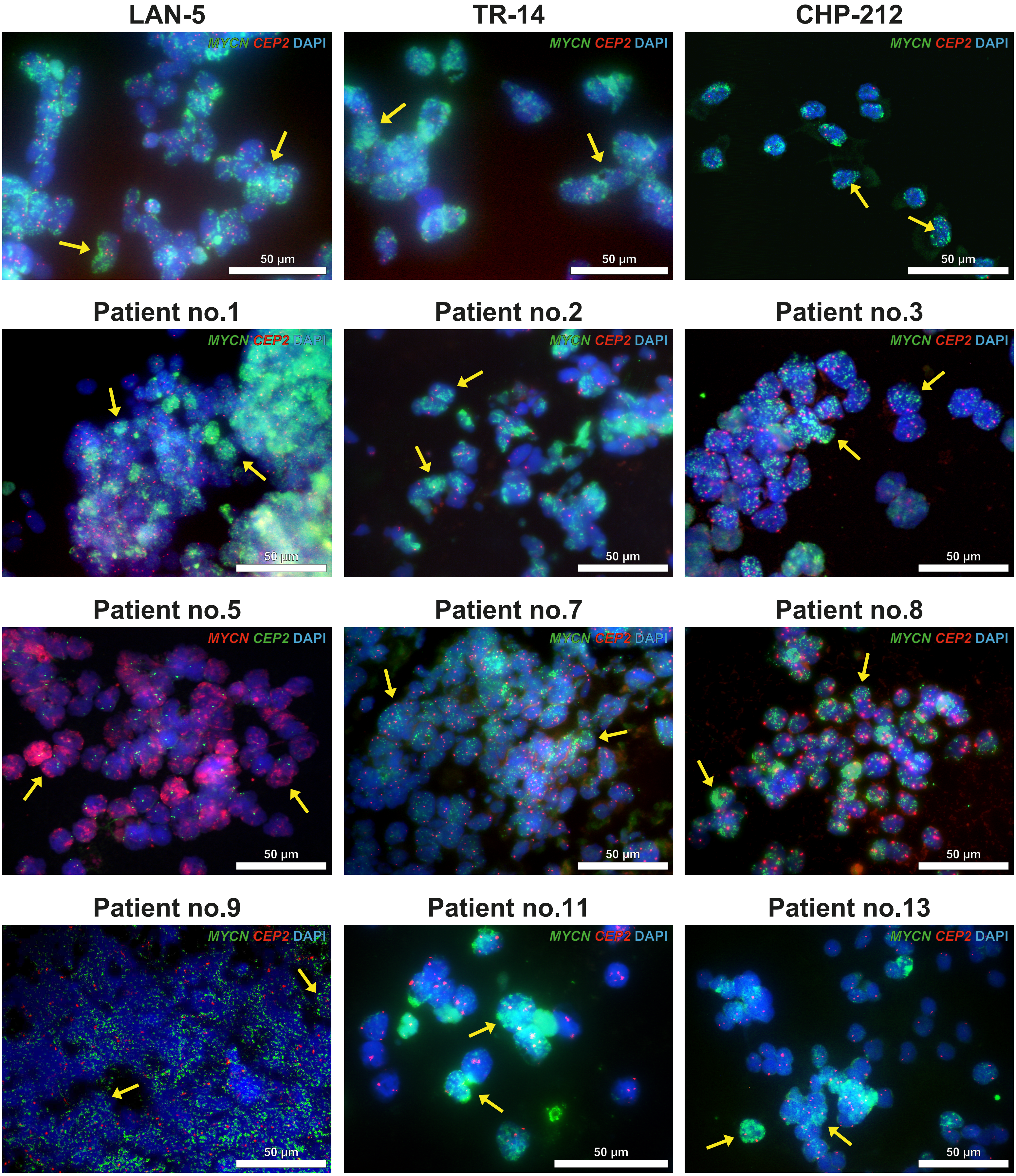

### SuppFigure2_NGS.pdf

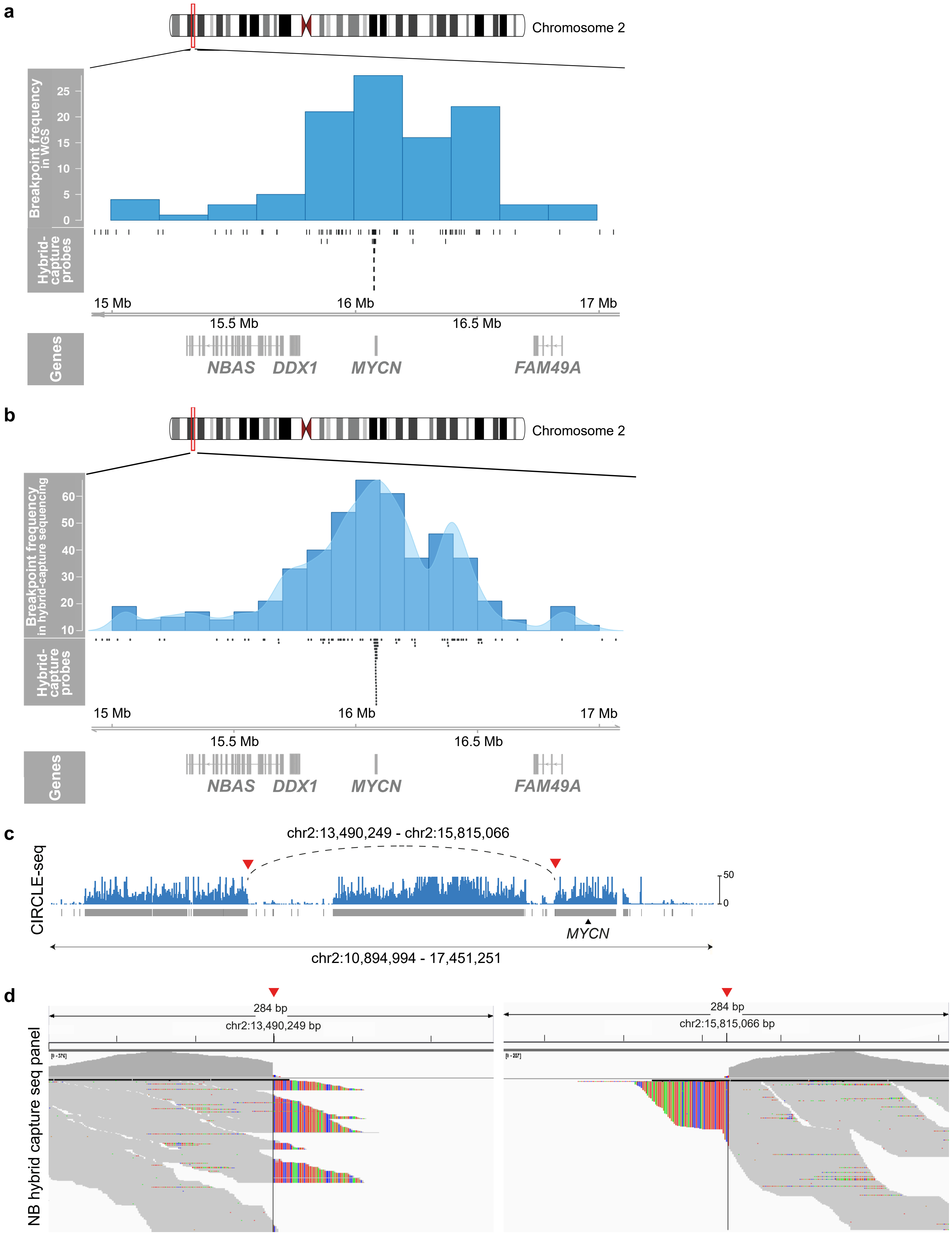

### SuppFigure5_sangercellLines.pdf

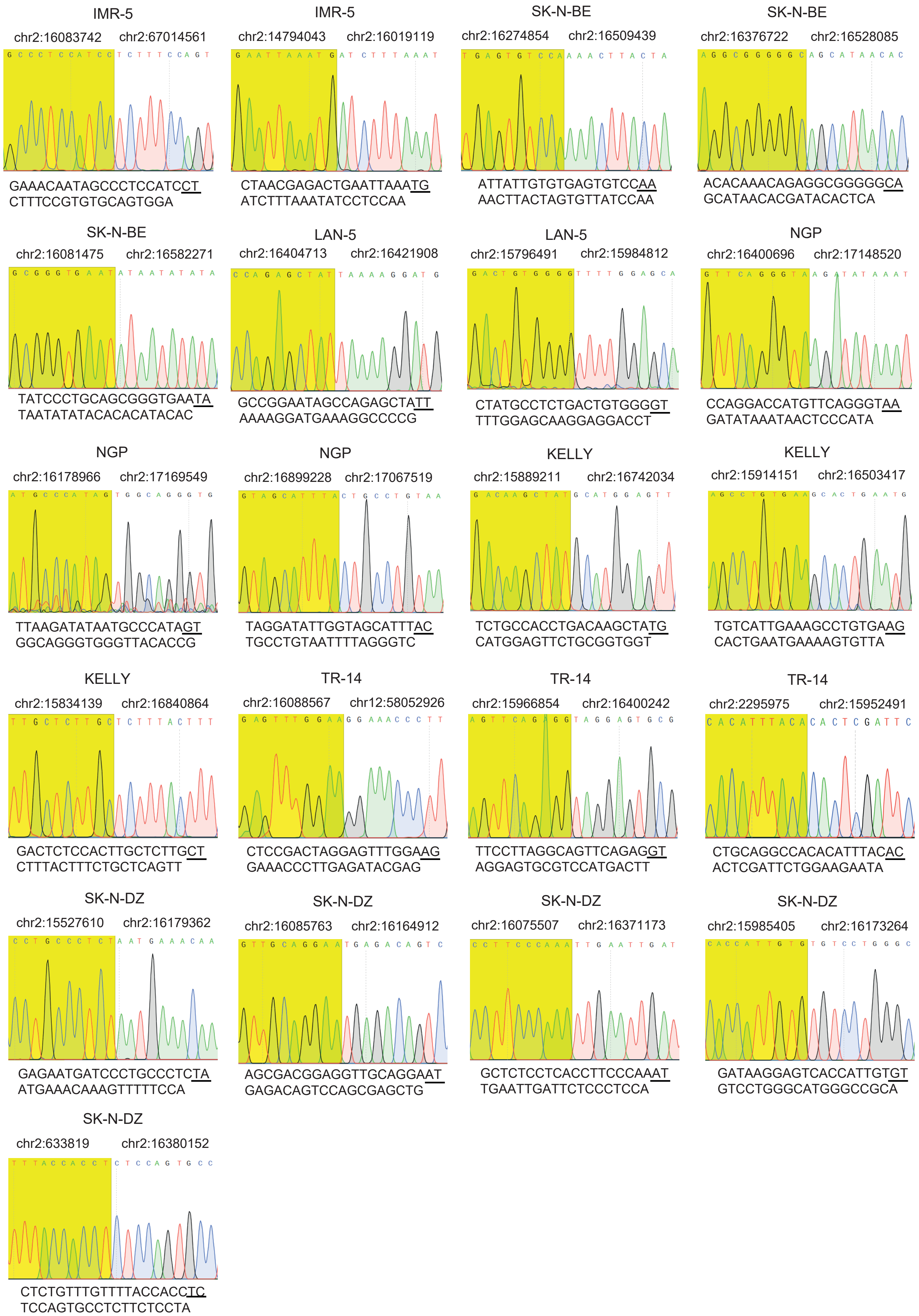

### SuppFigure6_sangerpatient.pdf

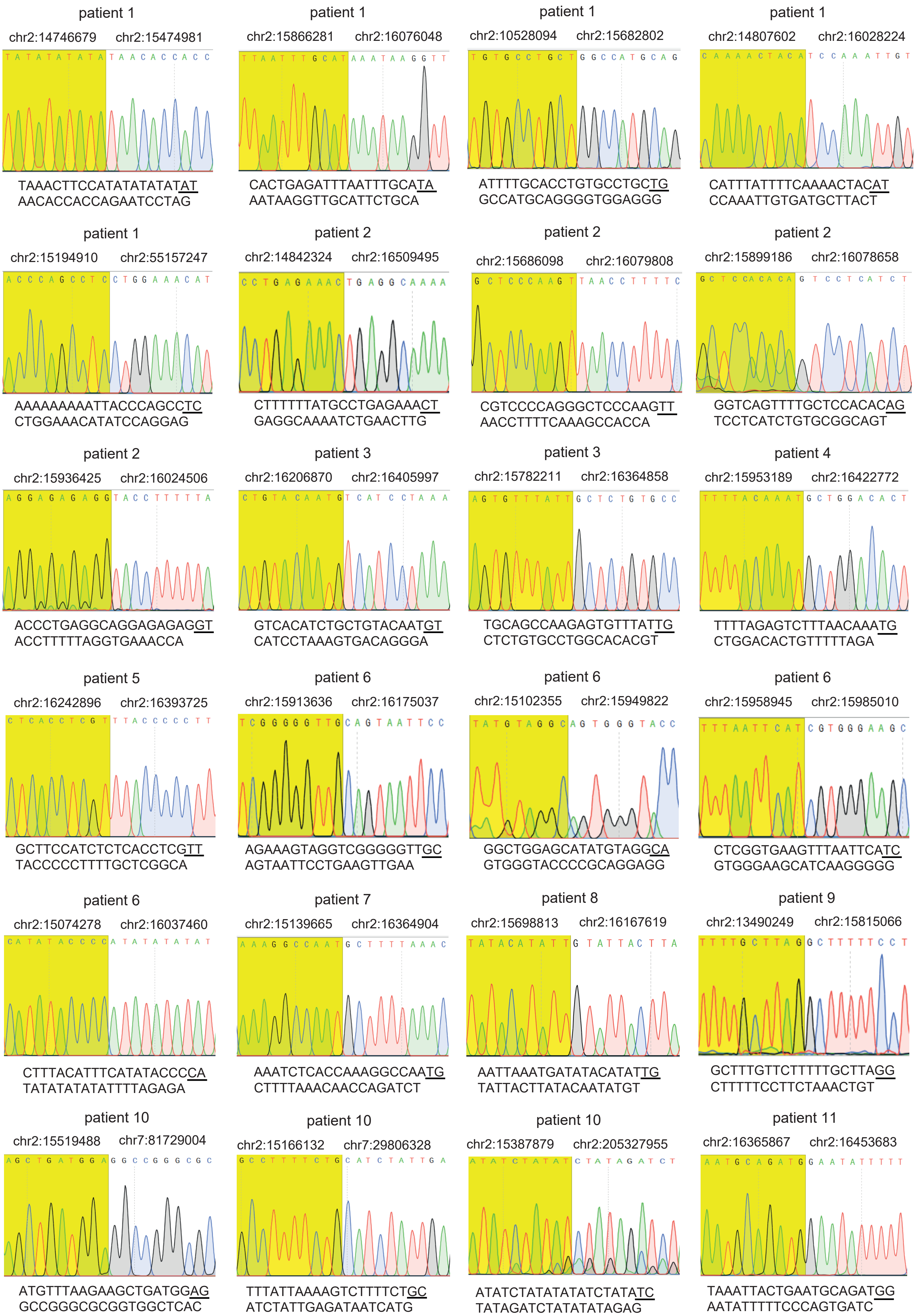

### SuppFigure7_additional_patients.pdf

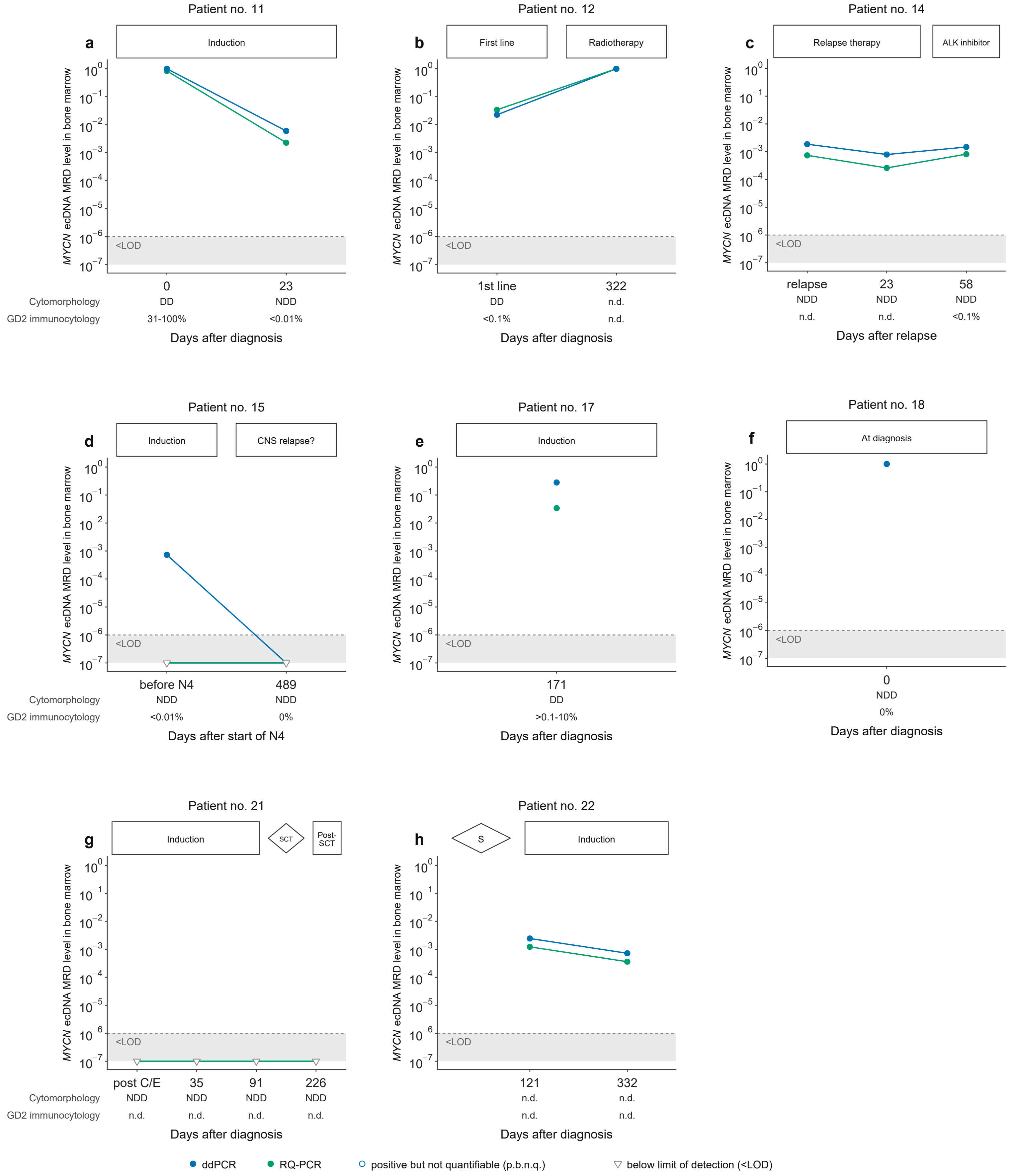
